# Host tropism determination by convergent evolution of immunological evasion in the Lyme disease system

**DOI:** 10.1101/2021.02.09.430532

**Authors:** Thomas M. Hart, Alan P. Dupuis, Danielle M. Tufts, Anna M. Blom, Simon Starkey, Ryan O. M. Rego, Sanjay Ram, Peter Kraiczy, Laura D. Kramer, Maria A. Diuk-Wasser, Sergios-Orestis Kolokotronis, Yi-Pin Lin

**Affiliations:** Division of Infectious Diseases, Wadsworth Center, New York State Department of Health, Albany, NY, USA; Department of Biological Sciences and State University of New York at Albany, NY, USA; Department of Biomedical Sciences, State University of New York at Albany, NY, USA; Department of Ecology, Evolution, and Environmental Biology, Columbia University, New York, NY USA; Division of Medical Protein Chemistry, Department of Translational Medicine, Lund University, Malmo, Sweden; Institute of Parasitology, Czech Academy of Sciences, České Budějovice, Czech Republic; Faculty of Science, University of South Bohemia, České Budějovice, Czech Republic; Division of Infectious Diseases and Immunology, University of Massachusetts Medical School, Worcester, MA, USA; Institute of Medical Microbiology and Infection Control, University Hospital of Frankfurt, Frankfurt, Germany; Department of Epidemiology and Biostatistics, School of Public Health, SUNY Downstate Health Sciences University, Brooklyn, NY, USA; Institute for Genomic Health, SUNY Downstate Health Sciences University, Brooklyn, NY, USA; Division of Infectious Diseases, Department of Medicine, College of Medicine, SUNY Downstate Health Sciences University, Brooklyn, NY, USA

**Keywords:** Lyme disease, CspA, Host tropism, *Borrelia*, Complement, Convergent evolution

## Abstract

Microparasites selectively adapt in some hosts, known as host tropism. Transmitted through ticks and carried mainly by mammals and birds, the Lyme disease (LD) bacterium is a well-suited model to study such tropism. LD bacteria species vary in host ranges through mechanisms eluding characterization. By feeding ticks infected with different LD bacteria species, utilizing feeding chambers and live mice and quail, we found species-level differences of bacterial transmission. These differences localize on the tick blood meal, and complement, a defense in vertebrate blood, and a bacterial polymorphic protein, CspA, which inactivates complement by binding to a host complement inhibitor, FH. CspA selectively confers bacterial transmission to vertebrates that produce FH capable of allele-specific recognition. Phylogenetic analyses revealed convergent evolution as the driver of such findings, which likely emerged during the last glacial maximum. Our results identify LD bacterial determinants of host tropism, defining an evolutionary mechanism that shapes host-microparasite associations.

## INTRODUCTION

The interactions between hosts and microparasites (e.g. bacteria, viruses, and protozoa) have arisen through numerous evolutionary events (1, 2), often resulting in generalist microparasites, which adapt to most host environments, or specialists, which selectively survive in particular host species. The association between microparasites and their respective hosts is defined as “host tropism (or host specialization)” (3). For vector-borne microparasites, such a host tropism can be dictated by not only host factors but also host constituents in the vectors (e.g. blood meals) (4). Because many bacteria varying in host specificity are involved in the infection cycle, the Lyme disease bacterium is one of the models regularly applied to investigate the host-microparasite interactions (4). Carried by *Ixodes* ticks, this disease is the most common vector-borne disease in the northern hemisphere (5). The causative agent of Lyme disease is a genospecies complex of the spirochete *Borrelia burgdorferi* sensu lato (also known as *Borreliella burgdorferi* sensu lato (NCBI taxid: 139), Lyme borreliae) (6). Among these genospecies, the most frequently isolated spirochetes from both ticks and vertebrate hosts are *B. afzelii*, *B. garinii*, and *B. burgdorferi* sensu stricto (hereafter referred to as *B. burgdorferi*) (7). Following the tick bite, spirochetes need to survive in tick blood meals, which permit transmission to the bite site of the host skin. Survival in host bloodstream then is a prerequisite for hematogenous dissemination and colonization of distant tissues, resulting in varied disease manifestations involving different organs in incidental hosts such as humans (8). In nature, Lyme borreliae can invade vertebrate reservoirs (mainly birds and rodents) but varying in their ability to infect these animals in a host-specific manner (9, 10). For example, some Lyme borreliae species, such as *B. burgdorferi*, can invade a wide range of hosts whereas others are selectively infectious in few host taxa (e.g. *B. afzelii* for rodents and *B. garinii* for birds). However, the molecular basis for such host tropism is largely unclear (9, 10).

The elimination of microparasites by host immune responses is a major bottleneck of infectivity, limiting the breadth of host competence (9, 11, 12). Complement is one of the first lines of host defenses in vertebrate animals and can be activated through three canonical routes: the classical, lectin, and alternative pathways (13). The activation of complement on the microparasite surface results in the formation of a protein complex called C3 convertase. C3 convertase is essential for complement function because it serves as a protease to cleave C3 protein to its active fragments. Further complement activation leads to the lysis of microparasites due to the formation of membrane attack complex pores (C5b-9) on microparasite surface. C3 convertase formed by alternative pathway activation, called C3bBb (composed of C3b and Bb proteins), activates more C3 molecules and results in a positive feedback of the C3 amplification loop, a unique characteristic of the alternative pathway. To avoid tissue damage from unwanted complement activation in the absence of microparasites, hosts possess complement inhibitors (13, 14). One of these inhibitors is factor H (FH), which binds to C3b and blocks further activation of complement (13, 15). Like other microparasites, Lyme borreliae exploit such host self-regulatory mechanisms by recruiting complement inhibitors on their surface (10, 16–18), which allows spirochetes to evade complement-mediated killing in the host bloodstream or tick blood meals (10, 16–18). In fact, spirochetes bind to FH through the production of several bacterial FH-binding proteins, called Complement Regulators Acquiring Surface Proteins (CRASPs): CspA, CspZ, and OspE-related proteins (19, 20). Among these CRASPs, CspA is uniquely required for bacterial transmission from nymphal ticks to vertebrate animals by binding to FH, resulting in complement evasion in those feeding ticks (21–23).

Interestingly, the ability of CspA to bind to FH from different vertebrate hosts varies by alleles and dictates the specificity of a Lyme borreliae species to survive in the sera of various animals (21, 24, 25). Specifically, the ability of CspA variants to bind to mammalian FH and survive in homologous sera correlates with the ability of these variants to promote tick-to-mouse transmission (21). These findings raise a possibility that CspA-mediated, FH binding-dependent complement evasion drives Lyme borreliae host tropism (9, 10). However, that possibility could not be fully demonstrated until we could elucidate the roles of CspA variants that do not promote tick-to-mouse transmission to confer tickborne transmission to other hosts. Additionally, if CspA is one of the spirochete determinants of host tropism, what evolutionary mechanisms give rise to the host-spirochete associations mediated by this protein?

In this study, we used Lyme disease spirochetes, avian and mammalian hosts, and the blood of mammals and birds as models to examine the role of complement in driving host tropism of microparasites. We further identified CspA as a molecular determinant for such tropism and elucidated the evolutionary mechanisms resulting in the allelically specific roles of this protein in conferring maintenance of microparasites in diverse hosts during the infection cycle.

## RESULTS

### Lyme borreliae genospecies differ in their levels of transmission to wild-type but not complement-deficient mice and quail

To examine tick-to-host transmission among Lyme borreliae species, we intradermally injected wild-type BALB/c (wild-type; WT) mice with *B. burgdorferi* B31-5A4, *B. garinii* ZQ1, or *B. afzelii* CB43. The tissues from B31-5A4- or CB43-infected mice had significantly greater spirochete burdens than those from uninfected mice (Fig. S1A to D). In contrast, except for the bladders from two mice, bacterial burdens in the tissues of ZQ1-infected mice were below detection limits (10 bacteria per 100ng total DNA) (Fig. S1A to D). To generate ticks harboring equal loads of spirochetes, we intradermally injected each of these strains into C3-deficient BALB/c mice (C3^−/−^) mice, which do not have functional complement. After allowing *I. scapularis* larvae to feed on these mice, we found similar burdens of these strains in all tested tissues, fed larvae, and post molting flat nymphs (Fig. S1 E to J).

We then permitted the nymphs carrying each of these strains to feed on WT mice and determined the spirochete burdens in the replete nymphs, the skin at the tick bite site, and blood from these animals at 7 days post feeding (dpf), and uninfected nymphs were included as control. Strains B31-5A4 and CB43 survived at these sites (~10^3^ spirochetes per tick or 10^2^ to 10^3^ spirochetes per 100ng DNA of tissues, Fig. 1A to C). Two out of six fed nymphs had undetectable loads of ZQ1 whereas the other four ticks have bacterial loads ranging from 39 to 111 spirochetes per tick (Fig. 1A). These low, variable values were not significantly different compared to uninfected nymphs (Fig. 1A, p = 0.52). Strain ZQ1 was also largely undetectable in tick bite sites and blood (three and five out of five bite sites and blood samples, respectively; Fig. 1B and C). Additionally, we fed nymphs carrying B31-5A4, CB43, or ZQ1 on C3^−/−^ mice and found that these strains were detected in fed nymphs, bite sites, and blood at similar levels (Fig. 1D to F). These results indicate that ZQ1 is less capable of surviving in fed nymphs and establishing infection than B31-5A4 and CB43 after ticks fed only on WT mice, but not on the mice lacking C3, suggesting that mouse complement dictates spirochete transmission. We further studied the tickborne transmission of these strains in a similar fashion using *Coturnix* quail, the avian model of Lyme disease (26, 27). We detected B31-5A4 and ZQ1 in fed nymphs, tick bite sites, and bloodstream (~10^4^ spirochetes per tick or 10^2^ to 10^3^ Lyme borreliae per 100ng DNA of tissues, Fig. 1G to I). Conversely, strain CB43 was not detected above the detection limit in these ticks and tissue samples (Fig. 1G to I). When the nymphs carrying each of these strains were allowed to feed on quail treated with *O. moubata* complement inhibitor (OmCI), which blocks quail complement at the level of activation of C5 (Fig. S2)(28), we found similar levels of spirochetes in fed nymphs, tick bite sites and blood (Fig. 1J to L). These data showed less efficient tick-to-quail transmission of CB43 than that of B31-5A4 and ZQ1, and as was the case in mice, quail complement dictates transmission efficiency among those spirochete strains.

**Figure 1.**
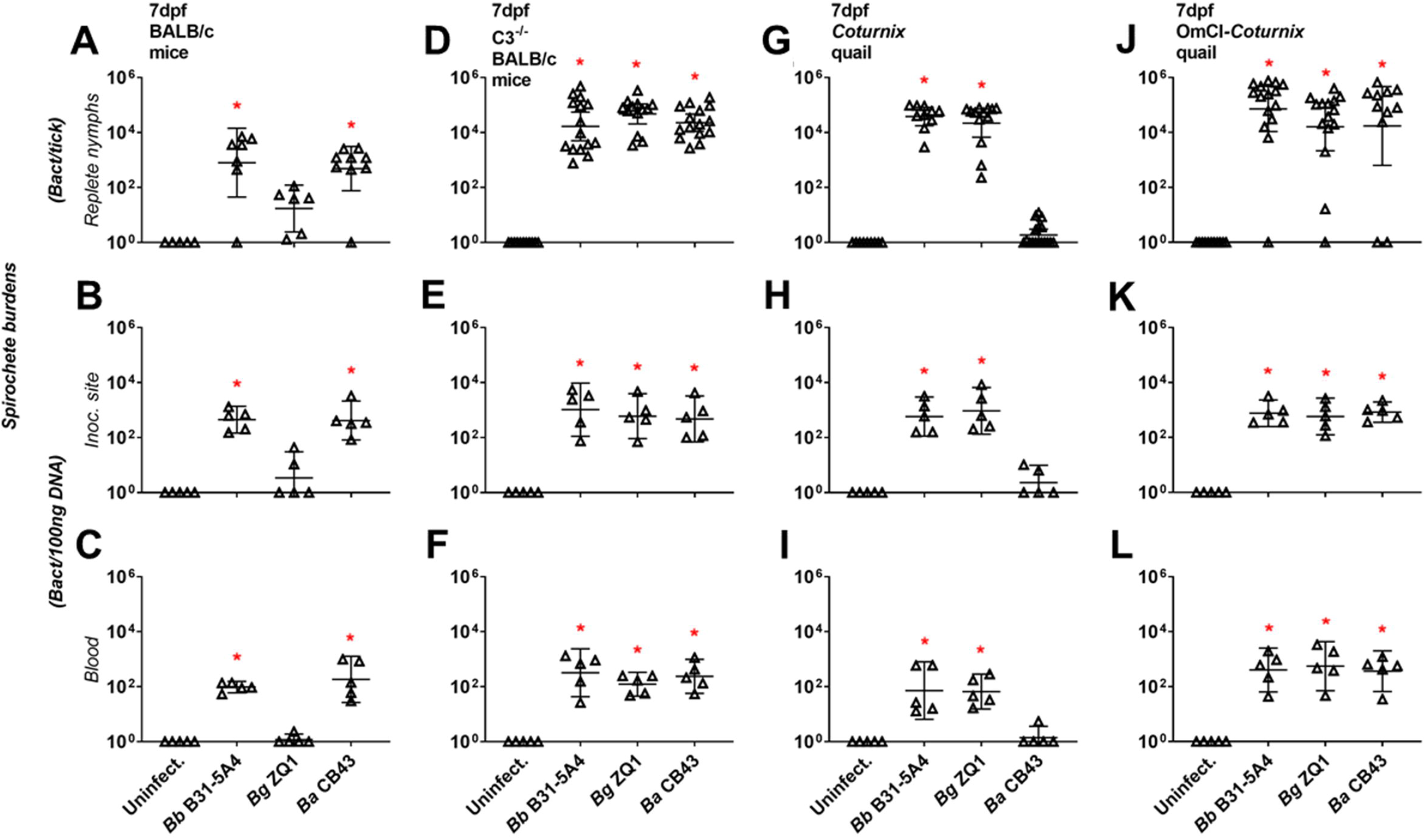
Lyme borreliae display species-level variation of tickborne transmission to wildtype but not complement-deficient mice and quail. *Ixodes scapularis* nymphs infected with *B. burgdorferi* B31-5A4 (“*Bb* B31-5A4”), *B. garinii* ZQ1 (“*Bg* ZQ1”), or *B. afzelii* CB43 (“*Ba* CB43”) fed on **(A-C)** BALB/c mice, **(D-F)** C3^−/−^ BALB/c mice, **(G-I)** *Coturnix* quail, or **(J-L)** OmCI-treated *Coturnix* quail. Uninfected nymphs and mouse and quail tissues were included as control (“Uninfect.”). Fed nymphs were collected upon repletion, and blood and the tick bite sites of skin were collected at 7 days post nymph feeding (“dpf”). Spirochete burdens in **(A, D, G, and J)** replete nymphs, **(B, E, H, and K)** tick bite site of skin (“Inoc. site”), and **(C, F, I, and L)** blood were determined by qPCR. For the burdens in tissue and blood samples, the resulting values were normalized to 100ng total DNA. Shown are the geometric means of bacterial loads ± 95% confidence interval of bacterial burdens in tissues and blood from 5 mice or quail per group or nymphs feeding on mice (7 nymphs carrying *Bb* B31-5A4, 6 nymphs carrying *Bg* ZQ1, or 9 nymphs carrying *Ba* CB43), C3^−/−^ mice (15 nymphs carrying *Bb* B31-5A4 or *Bg* ZQ1, or 13 nymphs carrying *Ba* CB43), quail (10 nymphs carrying *Bb* B31-5A4, 13 nymphs carrying *Bg* ZQ1, or 17 nymphs carrying *Ba* CB43), or OmCI-treated quail (15 nymphs carrying *Bb* B31-5A4, 11 nymphs carrying *Bg* ZQ1, or 15 nymphs carrying *Ba* CB43). Significant differences (P < 0.05, Kruskal-Wallis test with the two-stage step-up method of Benjamini, Krieger, and Yekutieli) in the spirochete burdens relative to uninfected ticks or tissues are indicated with an asterisk.

### The different abilities of Lyme borreliae genospecies to evade complement in tick blood meals determine mammalian- or avian-specific spirochete transmission

To examine the role that the source of the blood meals plays in determining spirochete transmission, nymphs infected with B31-5A4, ZQ1, or CB43 were allowed to feed on artificial feeding chambers with human blood, the mammalian blood representative (29)(Fig. 2A). Uninfected nymphs were included as controls. We found that B31-5A4 and CB43 survive in fed nymphs and blood (~10^3^ spirochetes per tick and 10^2^ spirochetes per 100 ng DNA of blood, Fig. 2B and C). In contrast, ZQ1 was undetectable in the majority of human blood-fed nymphs (six out of ten ticks) (Fig. 2B). Similarly, none of the blood samples fed on by ZQ1-infected nymphs had spirochete burdens significantly greater than uninfected blood (Fig. 2C). We also allowed nymphs carrying B31-5A4, ZQ1, or CB43 to feed on human blood treated with Cobra Venom Factor (CVF), which depletes human complement cascade from the level of C3 (30). We found similar burdens of these strains in the fed nymphs and the blood samples (~10^3^ spirochetes per tick and ~10^2^ spirochetes per 100 ng DNA of blood, Fig. 2D and E). These results indicate that ZQ1 is less competent than B31-5A4 or CB43 to survive in the tick blood meals from humans during transmission, and active human complement in the blood meals drives survival differences. We also performed similar work using quail blood and detected the strains B31-5A4 and ZQ1 in fed nymphs and blood (~10^3^ spirochetes per tick and ~10^2^ spirochetes per 100ng total DNA of blood, Fig. 2F and G). However, the strain CB43 was not detected in the majority of fed nymphs (7 out of 13 nymphs, Fig. 2F; P > 0.05 compared to uninfected nymphs). Further, spirochete burden values in blood samples fed on by nymphs carrying CB43 were indistinguishable from those of uninfected blood samples (Fig. 2G). Similar levels of each strain were observed when nymphs carrying each of these strains were fed on OmCI-treated quail blood (~10^3^ spirochetes per tick and 10^2^ spirochetes per 100 ng DNA of blood, Fig. 2H and I). These findings indicate that quail complement limits CB43 survival when ticks feed on quail blood.

**Figure 2.**
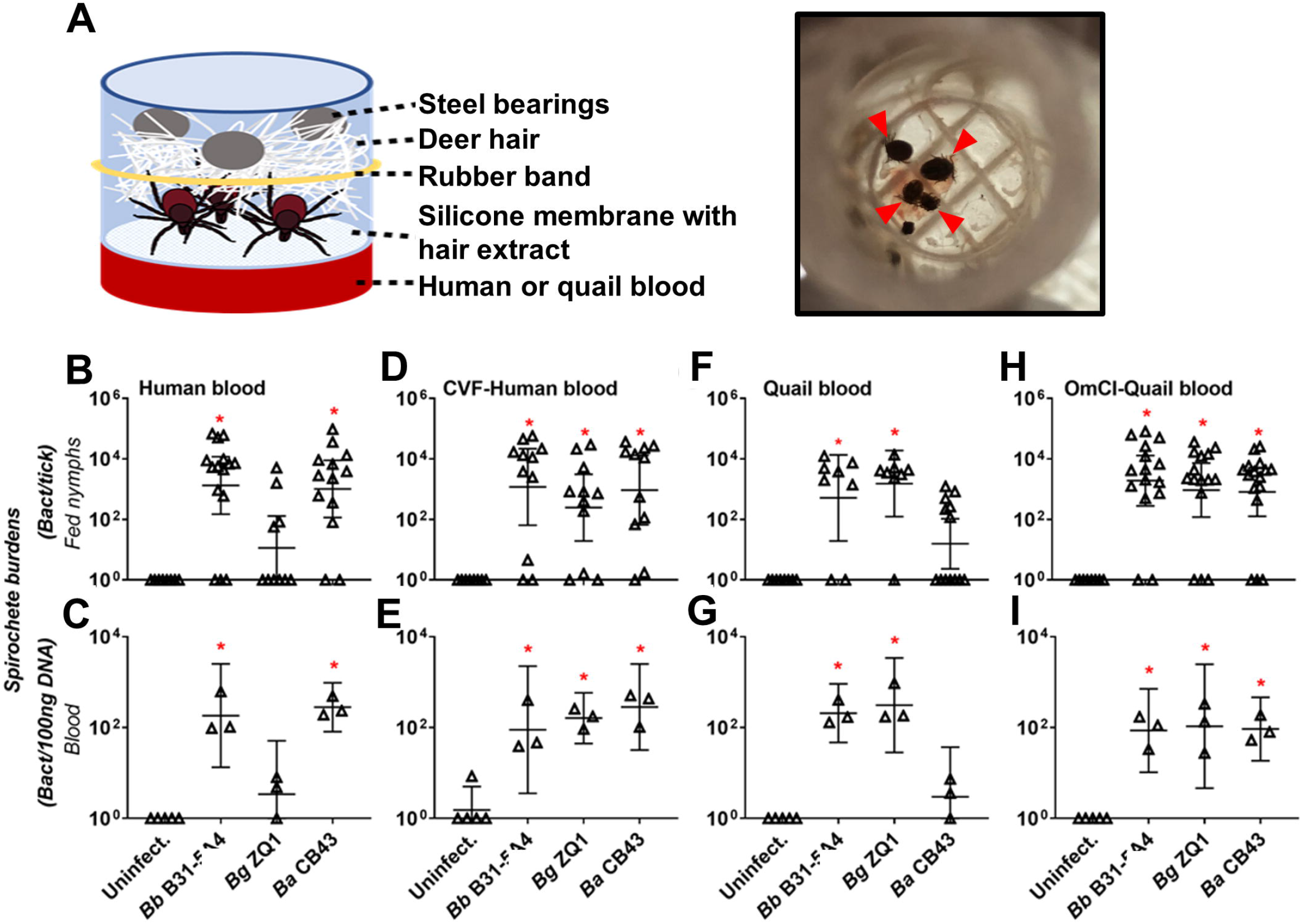
Complement in tick blood meals determines human or quail blood-specific spirochete transmission in feeding chamber. **(A) (left panel)** The schematic diagram showing the artificial feeding chamber that is used to examine the tickborne spirochete transmission in this study. **(Right panel)** The picture showing the engorged *I. scapularis* nymphs (indicated by arrows) in the chamber feeding on OmCI-treated quail blood. **(B-I)** *I. scapularis* nymphs infected with *B. burgdorferi* B31-5A4 (“*Bb* B31-5A4”), *B. garinii* ZQ1 (“*Bg* ZQ1”), or *B. afzelii* CB43 (“*Ba* CB43”), were allowed to feed in artificial feeding chambers submerging into six well plates containing **(B and C)** untreated or **(D and E)** CVF-treated human blood, or **(F and G)** untreated or **(H and I)** OmCI-treated quail blood. Blood was changed every 24-h and collected along with ticks on the fifth day of feeding. Uninfected nymphs and blood were included as control (“Uninfect.”). Spirochete burdens in **(B, D, F, and H)** fed nymphs and **(C, E, G, and I)** blood were determined by qPCR. The spirochete burdens in the blood were obtained by normalizing the resulting values to 100 ng total DNA. Shown are the geometric means of bacterial loads ± 95% confidence interval of bacterial burdens from the 3 human or quail blood samples or nymphs feeding on untreated human blood (15 nymphs carrying *Bb* B31-5A4, 10 nymphs carrying *Bg* ZQ1, or 13 nymphs carrying *Ba* CB43), CVF-treated human blood (11 nymphs carrying *Bb* B31-5A4, *Bg* ZQ1, or *Ba* CB43), quail blood (8 nymphs carrying *Bb* B31-5A4 or *Bg* ZQ1 or 13 nymphs carrying *Ba* CB43), or OmCI-treated quail blood (15 nymphs carrying *Bb* B31-5A4 or *Bg* ZQ1 or 16 nymphs carrying *Ba* CB43). Significant differences (P < 0.05, Kruskal-Wallis test with the two-stage step-up method of Benjamini, Krieger, and Yekutieli) in the spirochete burdens relative to uninfected ticks or blood are indicated with an asterisk

### CspA-mediated quail FH-binding activity promotes tick-to-quail transmission of spirochetes by complement evasion

We previously showed that a *cspA*-deficient mutant *B. burgdorferi* producing a spirochete outer surface protein, CspA, from the *B. burgdorferi* B31 (CspA_B31_) or *B. afzelii* PKo (CspA_PKo_) but not from *B. garinii* ZQ1 (CspA_ZQ1_), facilitates tick-to-mouse transmission by surviving in fed nymphs (21). This isogenic strain-specific transmission is dependent on the mouse FH-binding activity of these CspA variants to evade complement (21) (Table 1) and recapitulates the mouse-specific tickborne transmissibility of *B. burgdorferi* B31, *B. afzelii* CB43, and *B. garinii* ZQ1 (Fig. 1). (Note that CspA from *B. afzelii* CB43 (CspA_CB43_) shares 99% amino acid identity with CspA_PKo_, making these CspA variants likely confer similar FH-binding and transmission phenotypes (Fig. S3A)). To extend that isogenic strain-specific phenotype to other small mammals, we allowed nymphs carrying the that *cspA*-deficient *B. burgdorferi* producing CspA_B31_, CspA_PKo_, or CspA_ZQ1_ to feed on a rodent reservoir of Lyme borreliae, *Peromyscus leucopus*. We found that expression of CspA_B31_ and CspA_PKo_, but not CspA_ZQ1_, permitted spirochete transmission to that rodent species (Fig. S4). Further, our previous findings of CspA_ZQ1_ (and CspA_B31_) binding to quail FH and promoting survival in quail serum raised the hypothesis that CspA variants that bound FH drives tick-to-quail transmission (21) (Table 1). To test this hypothesis, quail were fed on by the nymphs carrying WT *B. burgdorferi* B31-5A15 (5A15), *B. burgdorferi* B31-5A4NP1Δ*cspA* harboring an empty vector (Δ*cspA*/Vector), or the *cspA*-deficient strain carrying plasmids to express CspA_B31_, CspA_ZQ1_, or CspA_PKo_ in the background of B31-5A4NP1Δ*cspA*. We also included an isogenic strain producing CspA_B31_-L246D, a CspA_B31_ mutant selectively devoid of quail FH-binding activity (21) (Table 1). Uninfected ticks were included as control. The strain 5A15 but not Δ*cspA*/Vector or *cspA*_B31_-L246D-complemented strain, had burdens above detection limits in the fed nymphs, tick bite sites and blood (Fig. 3A to C). Strains producing CspA_B31_ or CspA_ZQ1_ but not CspA_PKo_, had detectable burdens in the fed nymphs or tick bite sites and blood (~10^4^ spirochetes per tick and ~10^2^ spirochetes per 100 ng DNA of tissues or blood, Fig. 3A to C). In contrast, when nymphs carrying each of these spirochete strains were permitted to feed on OmCI-treated quail, all strains showed comparable burdens in fed nymphs, tick bite sites, and blood (Fig. 3D to F). These findings suggest that the CspA_ZQ1_ and CspA_B31_ as quail FH binders promote tick-to-quail transmission by evading complement, and CspA-mediated quail FH-binding activity dictates such a transmission.

**Table 1.**
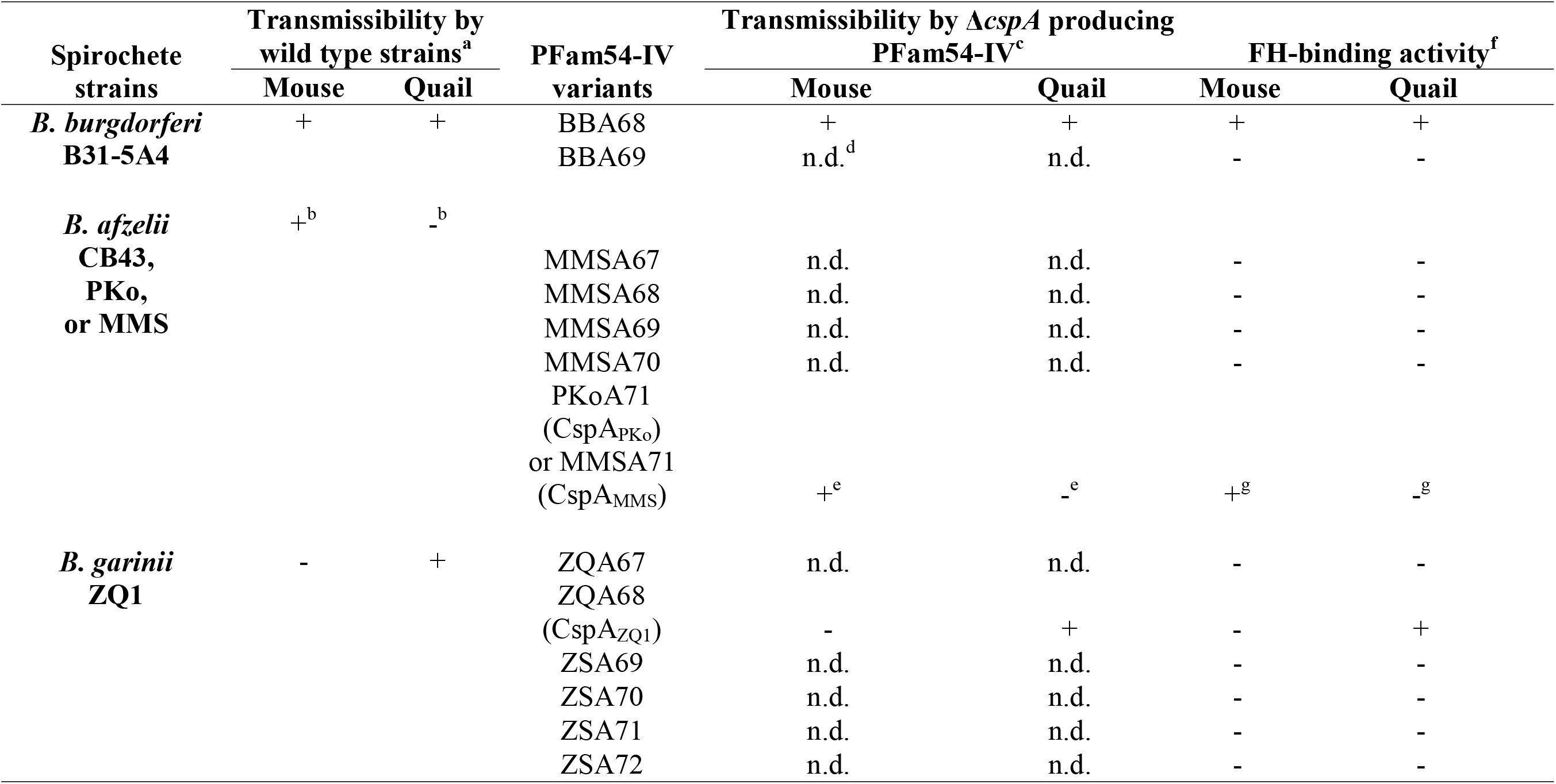
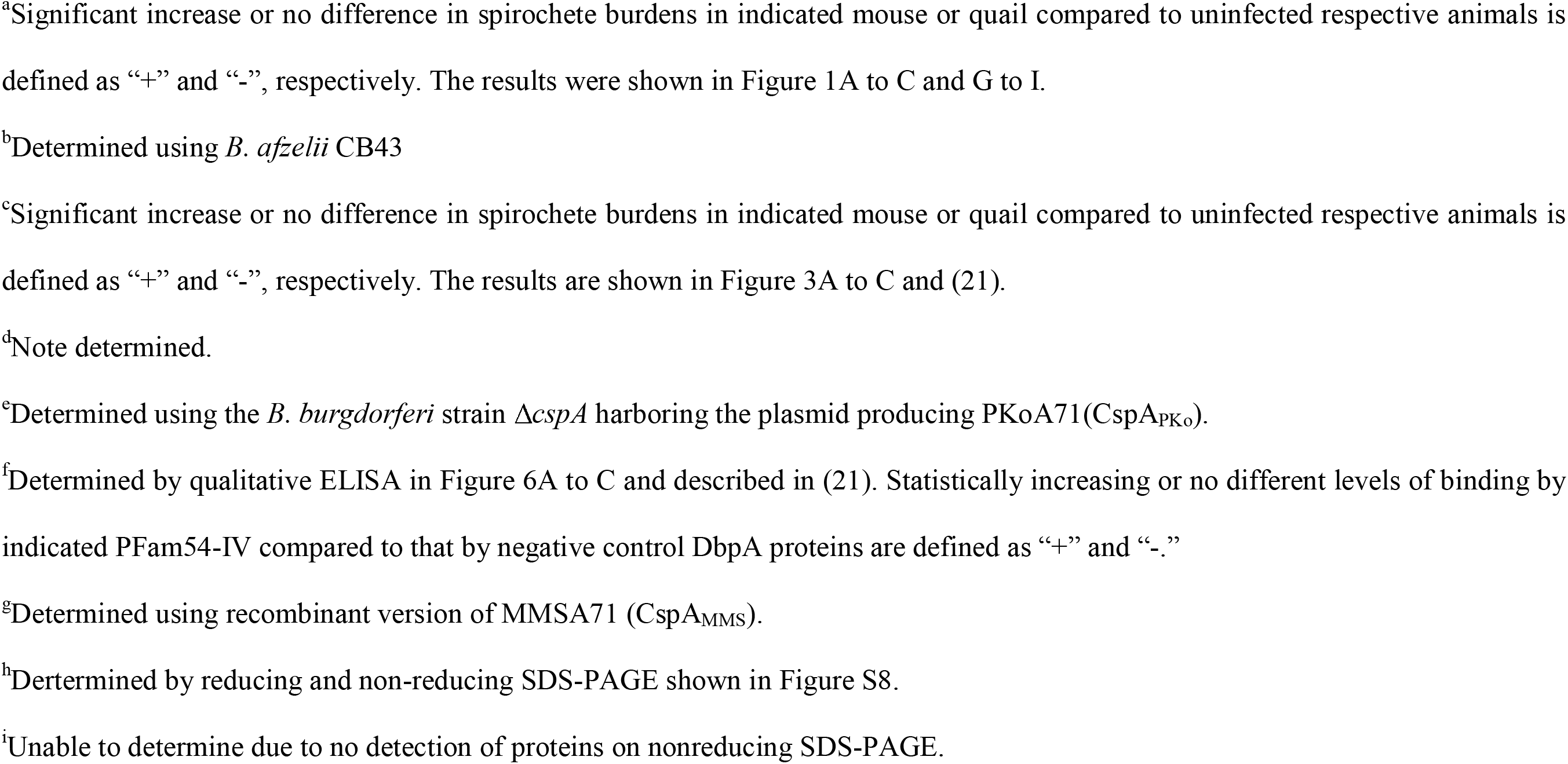
Summarized findings of allelically variable, host-specific transmissibility and FH-binding activity of *B. burgdorferi*, *B. afzelii*, and *B. garinii* and their derived PFam54-IV proteins in this study and the previous study (21).

| Spirochete strains | Transmissibility by wild type strains<sup>a</sup> | | PFam54-IV variants | Transmissibility by $\Delta cspA$ producing PFam54-IV<sup>c</sup> | | FH-binding activity<sup>f</sup> | |
| --- | --- | --- | --- | --- | --- | --- | --- |
|  | Mouse | Quail |  | Mouse | Quail | Mouse | Quail |
| *B. burgdorferi* B31-5A4 | + | + | BBA68<br>BBA69 | +<br>n.d.<sup>d</sup> | +<br>n.d. | +<br>- | +<br>- |
| *B. afzelii* CB43, PKo, or MMS | +<sup>b</sup> | -<sup>b</sup> | MMSA67<br>MMSA68<br>MMSA69<br>MMSA70<br>PKoA71<br>(CspA<sub>PKo</sub>)<br>or MMSA71<br>(CspA<sub>MMS</sub>) | n.d.<br>n.d.<br>n.d.<br>n.d.<br>+<sup>e</sup> | n.d.<br>n.d.<br>n.d.<br>n.d.<br>-<sup>e</sup> | -<br>-<br>-<br>-<br>+<sup>g</sup> | -<br>-<br>-<br>-<br>-<sup>g</sup> |
| *B. garinii* ZQ1 | - | + | ZQA67<br>ZQA68<br>(CspA<sub>ZQ1</sub>)<br>ZSA69<br>ZSA70<br>ZSA71<br>ZSA72 | n.d.<br>-<br>n.d.<br>n.d.<br>n.d.<br>n.d. | n.d.<br>+<br>n.d.<br>n.d.<br>n.d.<br>n.d. | -<br>-<br>-<br>-<br>-<br>- | -<br>+<br>-<br>-<br>-<br>- |

<sup>a</sup>Significant increase or no difference in spirochete burdens in indicated mouse or quail compared to uninfected respective animals is defined as “+” and “-”, respectively. The results were shown in Figure 1A to C and G to I.
<sup>b</sup>Determined using *B. afzelii* CB43
<sup>c</sup>Significant increase or no difference in spirochete burdens in indicated mouse or quail compared to uninfected respective animals is defined as “+” and “-”, respectively. The results are shown in Figure 3A to C and (21).
<sup>d</sup>Note determined.
<sup>e</sup>Determined using the *B. burgdorferi* strain $\Delta cspA$ harboring the plasmid producing PKoA71(CspA<sub>PKo</sub>).
<sup>f</sup>Determined by qualitative ELISA in Figure 6A to C and described in (21). Statistically increasing or no different levels of binding by indicated PFam54-IV compared to that by negative control DbpA proteins are defined as “+” and “-.”
<sup>g</sup>Determined using recombinant version of MMSA71 (CspA<sub>MMS</sub>).
<sup>h</sup>Dertermined by reducing and non-reducing SDS-PAGE shown in Figure S8.
<sup>i</sup>Unable to determine due to no detection of proteins on nonreducing SDS-PAGE.

**Figure 3.**
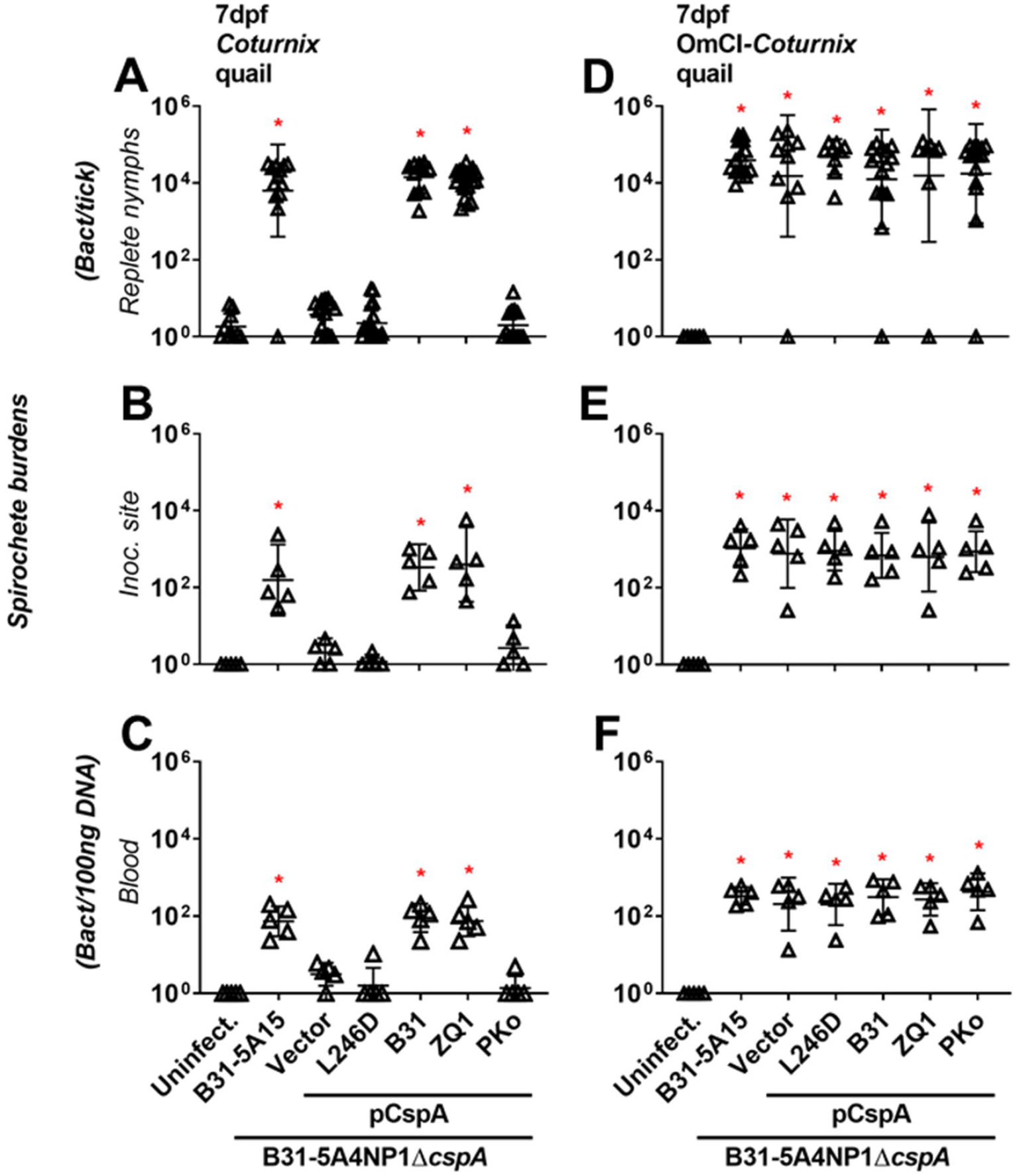
Polymorphic quail FH-binding activity of CspA confer the distinct transmissibility of Lyme borreliae to quail. *Ixodes scapularis* nymphs infected with WT *B. burgdorferi* B31-5A15 (“B31-5A15”), *B. burgdorferi* B31-5A4NP1Δ*cspA* transformed with an empty shuttle vector (“Vector”), or this deletion strain producing a mutated variant of CspA from *B. burgdorferi* B31 selectively devoid of quail FH binding activity (“L246D”), WT *B. burgdorferi* B31 (“B31”), *B. garinii* ZQ1 (“ZQ1”), or *B. afzelii* PKo (“PKo”) were allowed to feed on **(A-C)** untreated or **(D-F)** OmCI-treated quail. Uninfected nymphs and quail tissues were included as control (“Uninfect.”). Fed nymphs were collected upon repletion, and blood and tissues were collected at 7 days post nymph feeding (“dpf”). Spirochete burdens in **(A and D)** replete nymphs, **(B and E)** tick bite sites of skin (“Inoc. site”), and **(C and F)** blood were determined by qPCR. For the burdens in tissue samples, the resulting values were normalized to 100ng total DNA. Shown are the geometric means of bacterial loads ± 95% confidence interval of bacterial burdens from 5 quail tissues and blood or nymphs feeding on untreated quail (13 nymphs carrying the strains B31-5A15 or pCspA-PKo, 15 nymphs carrying the strains “Vector”, 12 nymphs carrying the strain pCspA-B31, 21 nymphs carrying the strain pCspA-ZQ1, or 17 nymphs carrying the strain pCspA-L246D), or the nymphs from OmCI-treated quail (15 nymphs carrying the strains B31-5A15, pCspA-B31, or pCspA-PKo, 8 nymphs carrying the strains “Vector”, 9 nymphs carrying the strain pCspA-L246D, 8 nymphs carrying the strain pCspA-ZQ1). Significant differences (P < 0.05, Kruskal-Wallis test with the two-stage step-up method of Benjamini, Krieger, and Yekutieli) in the spirochete burdens relative to uninfected ticks or quail tissues are indicated with an asterisk.

### Allelically variable, CspA-mediated FH-binding activity confers spirochete complement evasion in tick blood meals and transmissibility in a mammalian and avian blood-specific manner

We sought to examine whether CspA-mediated FH-binding activity facilitates spirochete evasion of complement in tick blood meals, and if that ability determines tickborne transmission in a host-specific manner. Human blood was allowed to be ingested by the nymphs carrying 5A15, Δ*cspA*/Vector, or this strain producing CspA_B31_, CspA_ZQ1_, CspA_PKo_, or CspA_B31_-L246D using feeding chambers. We detected 5A15, but not Δ*cspA*/Vector or CspA_B31_-L246D-producing strains, in fed nymphs and blood (Fig. 4A and B). The CspA_B31_- or CspA_PKo_-producing strains were found in fed nymphs and in human blood (~10^4^ spirochetes per tick (Fig. 4A) and more than 10 spirochetes per 100 ng total DNA of blood (Fig. 4B)). Conversely, the CspA_ZQ1_-producing strain was not detectable in these samples (Fig. 4A and B). When we allowed nymphs carrying the same strains to feed on CVF-treated human blood, all strains were detected at similar burdens in fed nymphs and blood (Fig. 4C and D). These results suggest that CspA_B31_ and CspA_PKo_, but not CspA_ZQ1_, permitted transmission to human blood by facilitating human FH-binding mediated complement evasion in tick blood meals. We also permitted the nymphs carrying the above-mentioned strains to feed on quail blood in the same fashion and found B31-5A15, but not Δ*cspA*/Vector or CspA_B31_-L246D, in the fed nymphs and blood had detectable burdens in both nymphs and blood (Fig. 4E and F). The CspA_B31_- or CspA_ZQ1_-producing strain was readily detected in fed nymphs and blood (Fig. 4E and F). Though three and two ticks carrying the CspA_PKo_ and CspA_B31_-L246D-producing strain, respectively, had burdens greater than detection limits, we did not detect any spirochetes in the remaining 10 nymphs (Fig. 4E). Additionally, the burdens of these strains in the blood were statistically indistinguishable from uninfected blood samples (Fig. 4F). When we performed similar experiments using OmCI-treated quail blood, all strains were found in nymphs and blood at comparable levels (Fig. 4G and H). These results show the contribution of quail FH-binding dependent complement evasion in tick blood meals in promoting transmission to quail blood, and CspA_B31_ and CspA_ZQ1_, but not CspA_PKo_, conferred these activities.

**Figure 4.**
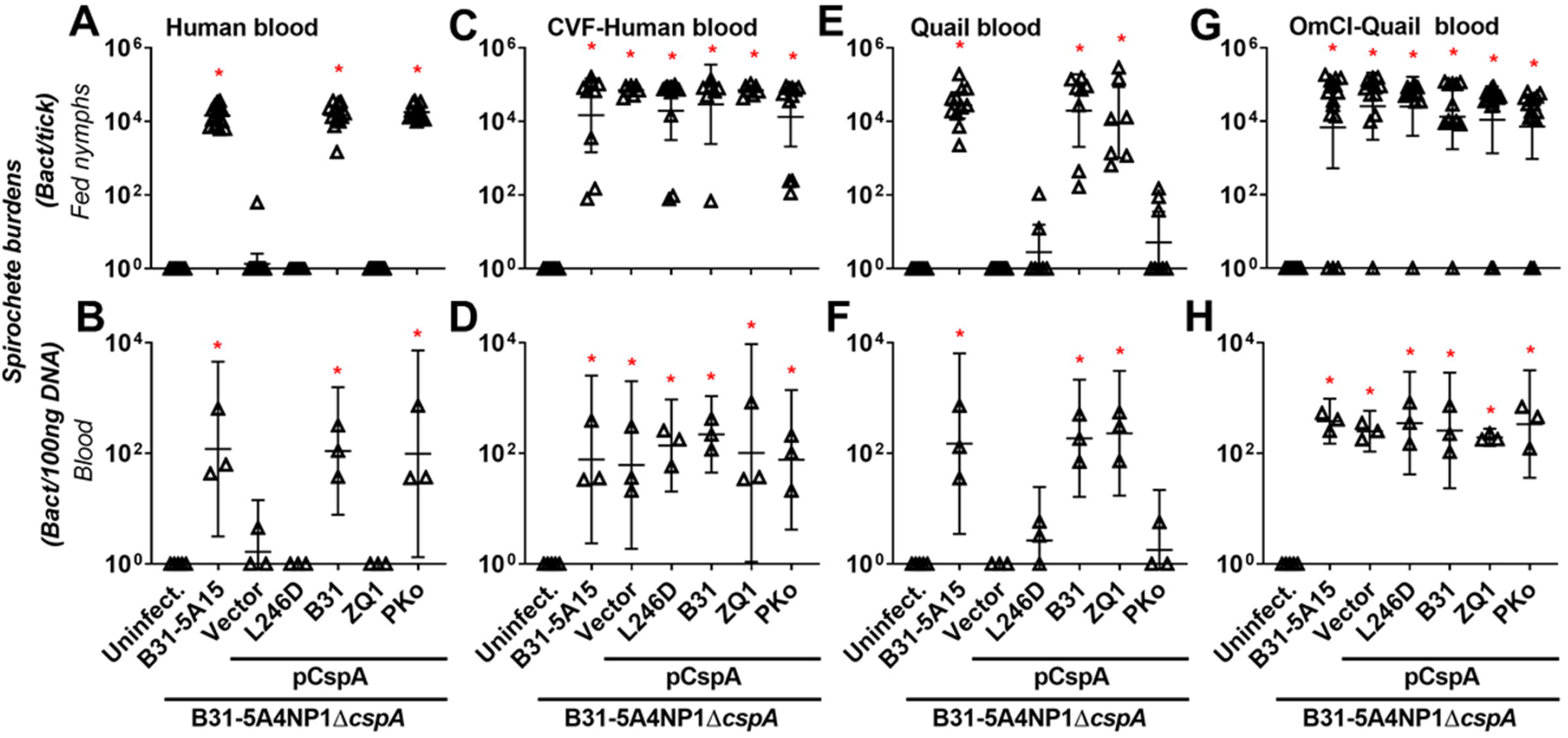
Allelically variable FH-binding activity of CspA dictates human- and quail blood-specific transmission in feeding chambers by evading complement in tick blood meals. *I. scapularis* nymphs infected with WT *B. burgdorferi* B31-5A15 (“B31-5A15”), *B. burgdorferi* B31-5A4NP1Δ*cspA* transformed with an empty shuttle vector (“Vector”), or this deletion strain transformed to produce a mutated variant of CspA from *B. burgdorferi* B31 selectively devoid of FH binding activity (“L246D”), WT *B. burgdorferi* B31 (“B31”), *B. garinii* ZQ1 (“ZQ1”), or *B. afzelii* PKo (“PKo”) were allowed to feed in feeding chambers submerged into 6-well plates containing **(A and B)** untreated or **(C and D)** CVF-treated human blood, or **(E and F)** untreated or **(G and H)** OmCI-treated quail blood. Blood was changed every 24-h and was collected along with ticks on the fifth day of feeding. Uninfected nymphs and blood were included as control (“Uninfect.”). Spirochete burdens from **(A, C, E and G)** fed nymphs and **(B, D, F, and H)** blood were determined by qPCR. For the burdens in tissue samples, the resulting values were normalized to 100ng total DNA. Shown are the geometric means of bacterial loads and 95% confidence interval of bacterial burdens from 3 human and quail blood samples, or nymphs feeding on untreated human blood (15 nymphs carrying the strains B31-5A15 or pCspA-B31, 14 nymphs carrying the strains “Vector” or pCspA-PKo, 17 nymphs carrying the strain pCspA-L246D, or 16 nymphs carrying the strain pCspA-ZQ1), or CVF-treated human blood (9 nymphs carrying the strain B31-5A15, 7 nymphs carrying the strains “Vector”, pCspA-B31, or pCspA-ZQ1, or 11 nymphs carrying the strains pCspA-L246D or pCspA-PKo), untreated quail blood (11 nymphs carrying the strains B31-5A15 or “Vector”, 8 nymphs carrying the strains pCspA-B31 or pCspA-PKo, or 7 nymphs carrying the strains pCspA-ZQ1 or pCspA-L246D) or OmCI-treated quail blood (15 nymphs carrying the strains B31-5A15, pCspA-PKo, or pCspA-ZQ1, 12 nymphs carrying the strains “Vector” or pCspA-B31, or 13 nymphs carrying the strain pCspA-L246D). Significant differences (P < 0.05, Kruskal-Wallis test with the two-stage step-up method of Benjamini, Krieger, and Yekutieli) in the spirochete burdens relative to uninfected ticks or quail tissues are indicated with an asterisk.

### CspA homologs showed discontinuous sequence variation and genospecies-specific polymorphisms

Given the finding that homologous CspA proteins from single strains of *B. burgdorferi*, *B. afzelii*, or *B. garinii* confer distinct host tropism, we examined CspA variation in publicly available sequences. CspA is nested in the fourth clade of a protein family encoded on the linear plasmid 54, lp54 (PFam54-IV) (41-81% nucleotide identity; Fig. 5A) (31, 32). However, the homology of all PFam54-IV proteins makes it difficult to easily identify CspA variants, leading to inaccurate annotations and misidentification (31). We thus compiled publicly available gene sequences encoding PFam54-IV available in GenBank from *B. burgdorferi*, *B. afzelii*, and *B. garinii*, and compared the pairwise nucleotide identities of codon alignments for these genes from each species. Within PFam54-IV-encoding genes of any one particular *B. burgdorferi* strain, we identified one-to-one orthologous genes based on sequence conservation (>95% identity, green in Fig. S5). In contrast, we found moderate conservation among PFam54-IV homologs lacking such one-to-one orthology (<81% identity, red and yellow in Fig. S5). Sequence divergence patterns (inlets in Fig. S5-S7) allowed us to identify genes encoding CspA_B31_, CspA_PKo_, and CspA_ZQ1_ as CspA orthologs in *B. burgdorferi*, *B. afzelii*, and *B. garinii*, respectively. Among these CspA orthologs, intraspecific diversity (i.e. within genospecies) exceeded 93% identity, while interspecific diversity (i.e. between genospecies) varied from 67 to 72% (Fig. 5B and C). These results suggest a genospecies-specific polymorphism among CspA variants, whereby variants of the same genospecies share notably high identity, while variants of different genospecies share relatively lower identities.

**Figure 5.**
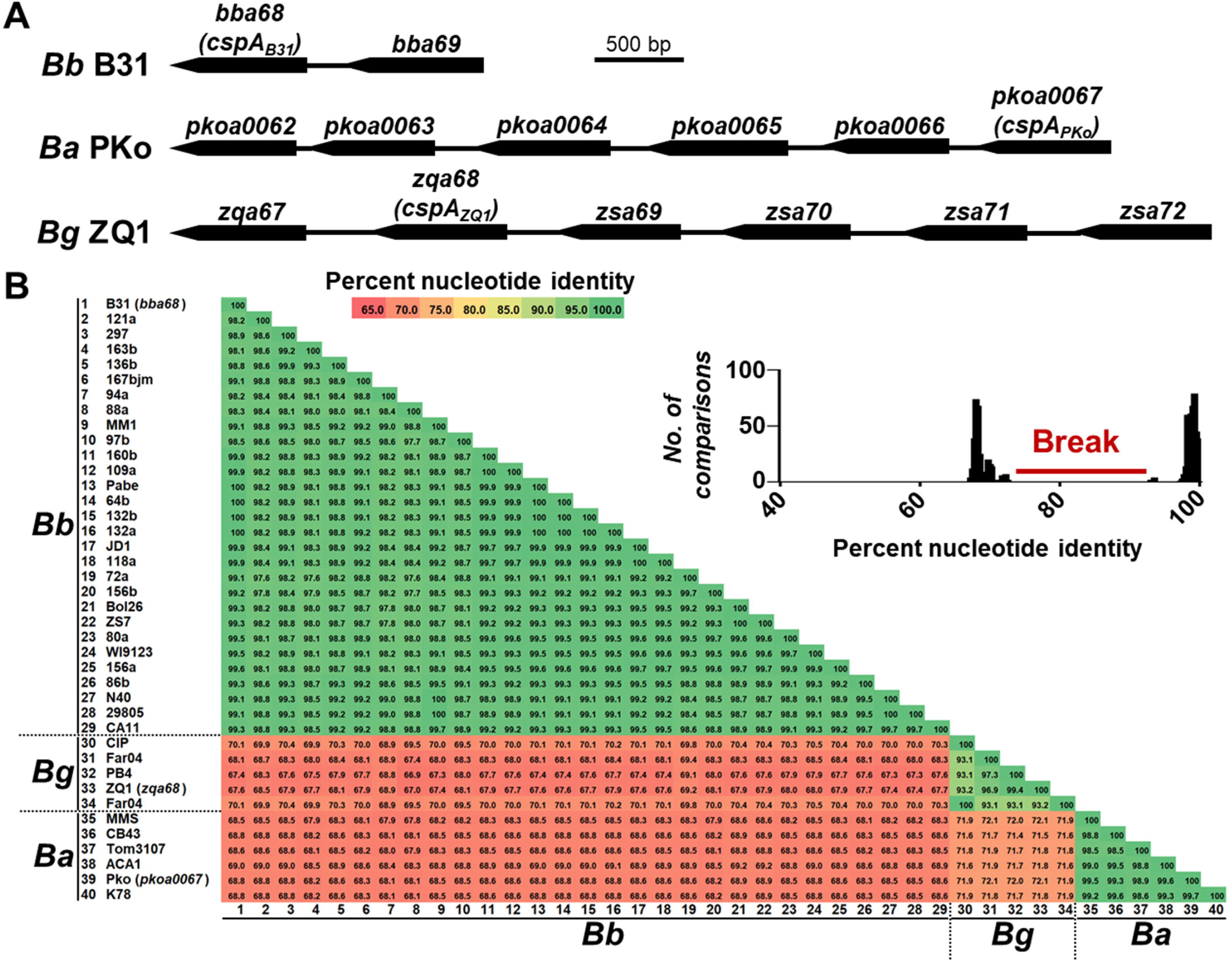
Pairwise comparisons reveal spirochete genospecies-specific CspA polymorphisms. **(A)** The synteny of Pfam54-IV proteins from *B. burgdorferi* B31, *B. afzelii* PKo, and *B. garinii* ZQ1 **(B)** CspA variants from *B. burgdorferi*, *B. afzelii*, and *B. garinii* identified in Fig. S5-S7 were aligned by codon using T-Coffee on the TranslatorX server. **(C)** The pairwise identity percentages were plotted versus the number of comparisons with those respective values, with bin widths of 0.25. Clear breaks in the pairwise sequence identity distribution were inspected to differentiate the highly identical (> 95%) from moderately divergent comparisons (< 80%). More than 95% identity in the comparisons of the variants between strains within spirochete genospecies but < 72% identity between genospecies indicate genospecies-specific *cspA* variation.

### Host-specific FH-binding activity of CspA variants arose through convergent evolution

An average identity of 74% among the genes encoding CspA and other Pfam54-IV proteins raises the possibility that non-CspA members of Pfam54-IV share FH-binding functions. We thus examined the mouse (*Mus musculus*) and quail FH-binding ability of Pfam54-IV from *B. burgdorferi* B31-5A4, *B. afzelii* MMS, and *B. garinii* ZQ1 using ELISA. Note that PFam54-IV members from *B. afzelii* MMS, PKo, and CB43 are nearly identical (>99% identity) (Fig. S3A and S6). We used Pfam54-IV of MMS to represent these proteins of *B. afzelii* given that the recombinant version of these proteins from MMS had been generated in our previous work (24). As expected, PFam54-IV from *B. burgdorferi* B31-5A4, *B. afzelii* MMS, and *B. garinii* ZQ1 did not bind to BSA (Fig. 6A). We found that CspA_B31_ and CspA_MMS_ bound to mouse FH at levels greater than a negative control spirochete protein, DbpA (21) (Fig. 6B). Despite a high concentration (2 μM) of other recombinant Pfam54-IV used, none bound to mouse FH over baseline levels seen with DbpA (Fig. 6B). Furthermore, we observed that CspA_B31_ and CspA_ZQ1_, but none of other tested Pfam54-IV, bound to quail FH (Fig. 6C). These results indicate that the host-specific and allelically variable FH-binding activity of Pfam54-IV is CspA-dependent.

**Figure 6.**
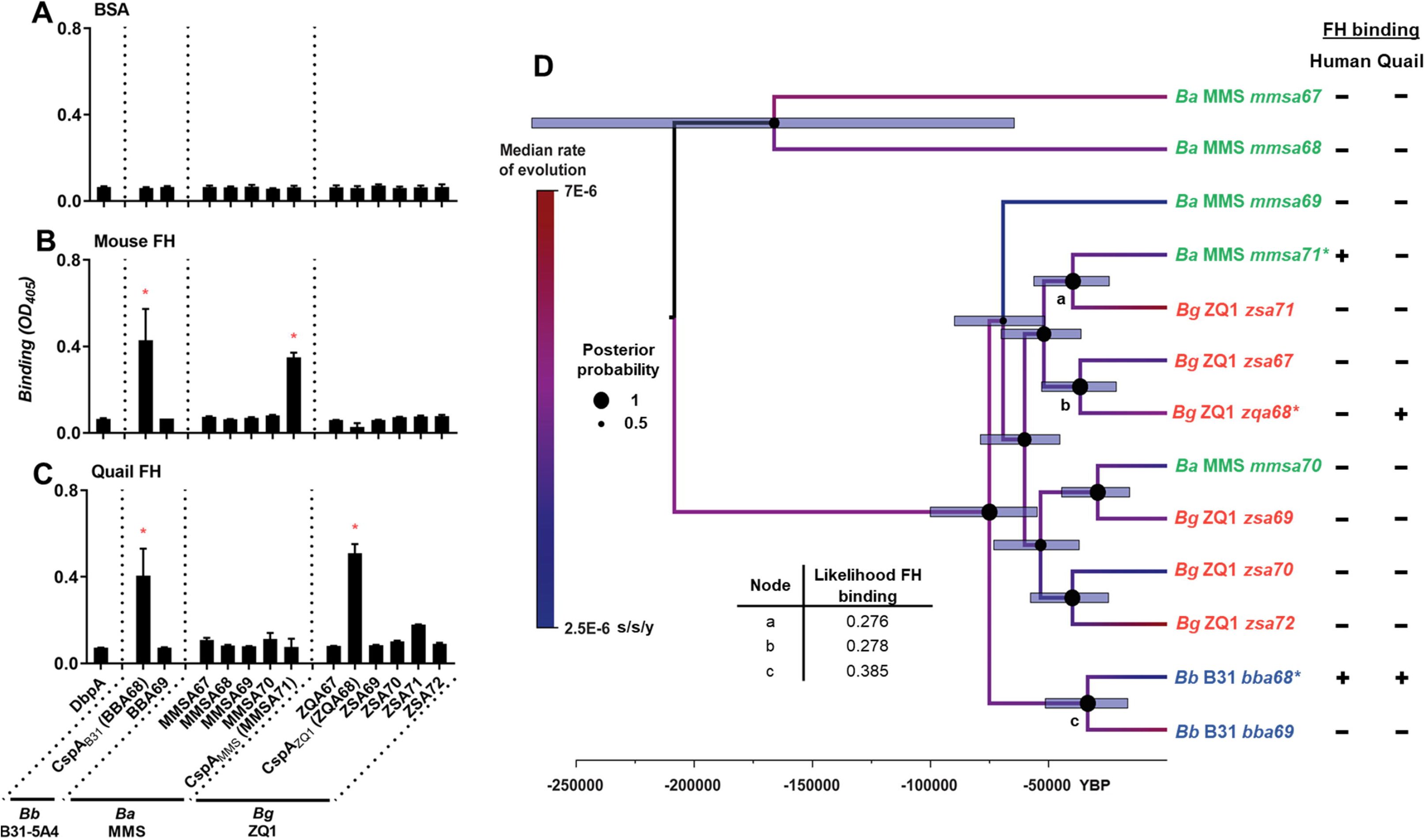
Allelically variable, host-specific FH-binding activity of CspA has emerged from convergent evolution. **(A to C)** 2 μM of histidine-tagged Pfam54-IV proteins from *B. burgdorferi* B31-5A4, *B. afzelii* PKo, or *B. garinii* ZQ1 or recombinant histidine-tagged DbpA from *B. burgdorferi* B31-5A4 (negative control) were added in triplicate to wells coated with of purified **(A)** BSA (negative control) or **(B)** mouse or **(C)** quail FH. The protein binding was measured by ELISA in three independent experiments. Each bar represents the geometric mean ± 95% confidence interval of three replicates in one representative experiment. Significant differences (P < 0.05, Kruskal-Wallis test with the two-stage step-up method of Benjamini, Krieger, and Yekutieli) in the levels of FH binding of indicated proteins relative to DbpA (‘‘*’’) are indicated. The ability of each protein in binding to factor H is summarized in Table 1. **(D)** A Bayesian phylogenetic reconstruction was generated based on the nucleotide sequences encoding Pfam54-IV proteins from *B. burgdorferi* B31 (blue), *B. afzelii* PKo (green), *B. garinii* ZQ1 (orange). In brief, these sequences were aligned by codon using T-Coffee on the TranslatorX webserver. Phylogenetic reconstruction was generated based on the resulting nucleotide alignment using BEAST2 with a lognormal uncorrelated relaxed clock with an estimated mutation rate of 4.75×10^−6^ substitutions/site/year (“s/s/y”) with a coalescent Bayesian skyline population. The resulting tree is drawn to scale, with branch lengths measured in the number of substitutions per site and rooted at the midpoint for clarity. The scale bar at the bottom represents an approximate timeline of evolution, in years before present (“YBP”), using the estimated substitution rate of 4.75×10^−6^ substitutions/site/year. Node bars represent the 95% highest posterior density of the node age. Node circles represent the posterior probability support. Branches are colored based on estimated the median substitution rate as per the legend to the left. **(E)** Maximum likelihood- and parsimony-based ancestral state reconstructions were used to predict FH-binding activities at ancestral nodes. FH-binding activities were not predicted at any node, and the likelihood of FH-binding activities at nodes immediately prior to CspA variants are indicated.

To further study the evolutionary mechanisms leading to host-specific and allelically variable FH-binding activity of CspA, we estimated phylogenetic relationships among gene sequences encoding PFam54-IV from *B. burgdorferi* B31, *B. afzelii* MMS, and *B. garinii* ZQ1. We found that those sequences of the same genospecies do not form monophyletic assemblages, but CspA variants grouped in separate clades with moderate to high internode branch support (Bayesian posterior probabilities (PP) of 0.81 at CspA_B31_ and CspA_MMS_ nodes and 0.86 at CspA_MMS_ and CspA_ZQ1_ nodes) (Fig. 6D). Similar branching patterns were seen when phylogenetic relationships were estimated among genes encoding PFam54-IV available on GenBank from *B. burgdorferi*, *B. afzelii*, and *B. garinii* (SH-aLRT/ultrafast bootstrap supports of 98.3/100% at CspA_*B. burgdorferi*_ and CspA_*B. afzelii*_ nodes and 84.4/74% at CspA_*B. afzelii*_ CspA_*B. garinii*_ nodes Fig. S8). We then tested the plausibility of this evolutionary scenario by placing CspA variants in the same clades (due to the same FH-binding functions, Fig. S9A, left panel) or cladding PFam54-IV variants from the same genospecies together (Fig. S9A, right panel). The results supported neither alternative phylogeny (Fig. S9B), in agreement with the phylogeny placing CspA variants in separated clades but not with every PFam54-IV protein from the same genospecies. Further, the supported phylogeny raises the possibility that CspA-mediated FH-binding activities arose from 1) a common FH-binding ancestor or 2) the convergent evolution of PFam54-IV (Fig. 6D). However, our results from maximum likelihood and parsimony-based tree-building methods, rejected the former possibility (Fig. 6D and E), indicating that the allelically variable, host-specific FH-binding activity of CspA is a result of convergent evolution within PFam54-IV. Based on a chromosome mutation rate estimated in a previous study (33), such an evolution event likely occurred approximately 15,000-55,000 years before present, coinciding with the end of the last glacial maximum (Fig. 6D).

## DISCUSSION

The constant interaction of microparasites and hosts allows the microparasites to adapt to each of the host environments, by which they can evolve to become specialists (4, 34). However, the fact that generalists are present in nature suggests that generalization of host ranges for those microparasites also confers fitness advantages (4, 34). For vector-borne microparasites, the process leading to host tropism can be driven by host-derived components (i.e. immune molecules or nutrients) either in the hosts or acquired by vectors (4). The molecular determinants and evolutionary mechanisms by which microparasites specialize or generalize to be associated with hosts are largely unclear. Reflected by the variable host tropism of different spirochete genospecies transmitted through *Ixodes* ticks, the Lyme disease bacterium is a well-suited model to study host-microparasite interactions (9, 10). *I. scapularis* ticks were shown in laboratory infections to carry *B. burgdorferi*, *B. afzelii*, and *B. garinii* at similar levels (35, 36), suggesting the use of this tick to represent *Ixodes* vectors of Lyme disease. Using a single tick species carrying each of the tested spirochete species allows for attribution of the observations solely to host and/or pathogen determinants, the emphasis of this study. Nonetheless, *B. afzelii* and *B. garinii* are not endemic to North America where *I. scapularis* are found, and are thus isolated from other *Ixodes* ticks (i.e. *I. ricinus* and *I. persulcatus*) in the field. *B. burgdorferi* is the only Lyme borrelia genospecies in this study that is circulated in *I. scapularis* in nature (5). Thus, utilizing *I. scapularis* as a vector representative may not completely address the role of vector competence in modulating host tropism of spirochetes (37, 38), which warrants further investigations. Additionally, Lyme borreliae-infected mice have commonly been used to generate nymphs harboring spirochetes through blood feeding by naïve larvae (39). However, rearing nymphs carrying similar burdens of each spirochete species in this fashion is difficult because wild-type mice do not maintain equal loads of these spirochetes (40, 41). Using complement-deficient mice (C3^−/−^ mice), we found similar burdens of *B. burgdorferi* B31-5A4, *B. afzelii* CB43, and *B. garinii* ZQ1 in fed larvae, post-molting flat nymphs, and the tissues derived from spirochete-infected mice. This result provides a strategy to overcome the difficulty in tick-rearing and infection, and supports the concept that complement controls the spirochete infectivity during infection (21, 40, 42).

No definitive studies have been performed to test the long-held model that complement evasion by spirochetes determines Lyme borreliae host tropism (43). A hurdle for such an investigation is the inability to easily maintain and/or persistently infect non-mammalian hosts, such as birds (44–54). Though some wild-birds have been brought into laboratories to study spirochete infectivity (44–54), molecular mechanisms have not been elucidated because of the lack of avian-specific reagents. We and others have intradermally inoculated Lyme borreliae into *Coturnix* quail as this domestic bird can sustain detectable spirochete burdens for more than eight weeks (26, 27). We thus allowed ticks carrying spirochetes to feed on quail, similar to previous work performed in this species and other domestic aves (55–57). We found that *B. garinii* ZQ1 and *B. burgdorferi* B31-5A4 survive in fed ticks and are transmitted to quail whereas *B. afzelii* CB43 did not. These results demonstrate that the genospecies variation of spirochete transmissibility to birds, in agreement with prior studies [reviewed in (9)], supporting the use of quail as an avian host representative. In contrast, when nymph feeding was performed on wild-type mice, *B. afzelii* CB43 and *B. burgdorferi* B31-5A4 survived in fed ticks and migrated to these animals while *B. garinii* ZQ1 did not. All three species survived in fed ticks and are transmitted to complement-deficient quail or mice. In support of previous *in vitro* evidence (43, 58), this study establishes complement evasion by spirochetes as a driver of Lyme borreliae host tropism. As the presence of complement in vertebrate blood, our findings raise a possibility that spirochetes must evade host complement specifically in the blood meal. However, tick feeding on live animals may introduce confounding factors of blood meal-independent, complement-mediated clearance (59). Using different sources of blood in “artificial feeding chambers” without the involvement of animals allows us to demonstrate that spirochete evasion to complement in blood meals dictates Lyme borreliae host tropism (29).

Both *Ixodes* ticks and Lyme borreliae produce complement-inactivating proteins to facilitate feeding and pathogen transmission (8, 60–62). Supported by many Lyme borreliae proteins are polymorphic, one attractive hypothesis is that polymorphisms in these proteins contribute to host tropism. Regrettably, using wild-type spirochete strains may not delineate the contribution these proteins individually because these spirochetes generate multiple polymorphic complement-inactivating proteins during transmission (22, 63–66). Thus, identical spirochete background strains have been used to express genes or alleles (also known as “isogenic strains”) to define bacterial determinants of particular phenotypes (21) (67, 68). We have previously shown that isogenic strains in a *cspA*-deficient background producing CspA from *B. burgdorferi* B31 or *B. afzelii* PKo, but not *B. garinii* ZQ1, promote tick-to-mouse transmission. The ability of ticks to transmit Lyme borreliae is contingent on the ability of the spirochete to survive in fed ticks, consistent with the phenotypes observed using wild-type genospecies (21). Such differential transmissibility depends on the ability of CspA to bind to mammal FH in the presence of mouse complement, leading to the possibility that allelically-variable, CspA-mediated FH-binding activity dictates host tropism (21, 24, 25). Nonetheless, that concept requires identification of the animals that are susceptible to Lyme borreliae that express ZQ1-derived CspA. In agreement with our previous work showing CspA from ZQ1 and B31 but not PKo binds to quail FH (21), we found the ZQ1- and B31- (but not PKo-) derived CspA facilitates tick-to-quail transmission of spirochetes. We further showed that such an allele-dependent, quail-specific transmission is determined by the presence of quail complement and the ability of CspA to bind to this species’ FH. Our results using feeding chambers, isogenic bacterial strains and complement-intact or - deficient human or quail blood demonstrated that CspA is a determinant of Lyme borreliae host tropism by promoting host-specific, FH-binding-dependent complement evasion in tick blood meals.

CspA, like other Pfam54-IV proteins, experienced numerous events of duplications and deletions, resulting in moderate identity (~40-80%) among the variants in this protein family (31, 32, 69). These observations suggest rapid evolution, potentially indicating novel functions in this region (31, 32, 70). This notion is supported by some important functions (e.g. complement evasion, cell adhesion, plasminogen binding, and tissue colonization) identified for CspA and other Pfam54 proteins (21, 71–74). However, the moderate sequence similarity from limited number of strains in those studies makes it difficult to definitively determine the one-to-one orthologs of each Pfam54-IV member (31, 32), creating hurdles to further investigate the evolutionary mechanisms giving rise to those functions. By comparing the sequences that encode PFam54-IV from different strains within each genospecies, we found that some comparisons showed moderate identity (<80%), while others shared a high degree of identity (~90% or more).

These results allow us to define these highly similar sequences as paralogs. These findings also suggest that the phenotypes conferred by one Pfam54-IV protein (e.g. host-specific FH-binding activity/transmissibility) are likely shared by its orthologs from different strains within the same genospecies (75). Additionally, we observed notably less identity (<79%) when comparing genes encoding Pfam54-IV among different spirochete genospecies, indicating genospecies-specific polymorphisms. These results, combined with the fact that CspA variants confer variable complement evasion and host tropism (21, 24, 25), are similar to findings of allelically-variable phenotypes in other Lyme borreliae polymorphic proteins (65, 67, 68). Further, these data suggest that any unidentified functions of CspA or other Pfam54-IV may vary among the strains from different genospecies.

Some phenotypes are shared among multiple Pfam54-IV proteins from the same Lyme borreliae genospecies (e.g. C7 and C9 binding-mediated complement evasion by Bga66 and Bga72 from *B. bavariensis*) (71). This finding raises the possibility that FH-binding is a common feature for non-CspA Pfam54-IV proteins within the same genospecies. However, our finding showing no detectable mouse or quail FH-binding to non-CspA Pfam54-IV indicates that the FH-binding activity is unique to CspA, in agreement with the results of human FH-binding activity of Pfam54-IV (24). Further, our data from phylogenetic reconstructions rejected the possibility of functional cladding of CspA or a common FH-binding ancestor, but supported convergent evolution as the mechanism leading to such an allelically variable, CspA-mediated FH-binding activity (76). With the fact that such an activity dictates Lyme borreliae host tropism, allelically variable, CspA-mediated FH-binding activity would effectively isolate different populations of Lyme borreliae in their respective hosts in nature, resulting in adaptive radiation of spirochetes. An intriguing question is whether such adaptation is the result of allopatric ecological speciation of the Lyme borreliae (77–82). However, the phylogenetic reconstruction of Pfam54-IV suggests that CspA of *B. burgdorferi* evolved approximately 15,000 to 55,000 years ago. The emergence of the CspA variants thus likely occurred after ancient Lyme borreliae speciation as the earliest common ancestor of *B. burgdorferi* in North America was dated at ~ 60,000 years ago (33). Rather, that timeline of CspA emergence (~15,000-55,000 years ago) coincides with the latter half of the last glacial maximum in North America and Europe (83, 84). An attractive possibility can be considered that massive climatic changes triggered ecological shifts necessitating new strategies to be maintained in the enzootic cycle, such as allelically variable, CspA-mediated FH-binding activity. Using the Lyme disease bacterium as a model, this work is a pioneering study defining the mechanisms that dictate host tropism of microparasites, identifying the molecular determinants, and elucidating the evolutionary drivers of such host-microparasite associations. These findings will provide significant impacts into the origin of a vector-borne enzootic cycle and establish the groundwork for future studies to investigate the mechanisms in shaping host-microparasite interaction.

## MATERIALS AND METHODS

### Ethics statement

All mouse and quail experiments were performed in strict accordance with all provisions of the Animal Welfare Act, the Guide for the Care and Use of Laboratory Animals, and the PHS Policy on Humane Care and Use of Laboratory Animals. The protocol was approved by the Institutional Animal Care and Use Committee (IACUC) of Wadsworth Center, New York State Department of Health (Protocol docket number 19-451) and Columbia University (Protocol docket number AC-AAAO4551). All efforts were made to minimize animal suffering.

### Mouse, quail, tick, bacterial strains, OmCI, and FH

BALB/c mice were purchased from Taconic (Hudson, NY). C3^−/−^ mice in BALB/c background were generated from the C3^−/−^ (C57BL/6) purchased from Jackson Laboratory (Bar Harbor, ME) as described in our previous study (21). *P. leucopus* mice were ordered from *Peromyscu*s genetic stock center at University of South Carolina (Columbia, SC). *Ixodes scapularis* tick larvae were purchased from National Tick Research and Education Center, Oklahoma State University (Stillwater, OK) or obtained from BEI Resources (Manassas, VA). Lyme borreliae-infected nymphs were generated as described in the section “Generation of ticks carrying Lyme borreliae.” The *Borrelia* and *Escherichia coli* strains used in this study are described in Table S1. *E. coli* strains DH5α, M15, and derivatives were grown in Luria-Bertani (BD Bioscience) broth or agar, supplemented with kanamycin (50 μg/ml), ampicillin (100 μg/ml), or no antibiotics as appropriate. All *B. burgdorferi*, *B. afzelii*, and *B. garinii* strains were grown in BSK-II completed medium supplemented with kanamycin (200 μg/mL), streptomycin (50 μg/mL), gentamicin (50 μg/mL), or no antibiotics (see Table S1). Mouse FH was purchased from MyBiosource. Quail FH and recombinant OmCI proteins were generated as described previously (21, 26, 28).

### Mouse infection using needle inoculation

Four-week-old female BALB/c or C3^−/−^ mice in BALB/c background were used for experiments involved in needle infection of Lyme borreliae strains. Mice were infected by intradermal injection as previously described (68) with 10^6^ of *B. afzelii* CB43, *B. garinii* ZQ1, or *B. burgdorferi* B31-5A4. The plasmid profile of the strain B31-5A4 was verified prior to infection as described to ensure the stability of the vector and no loss of plasmids (85, 86). As the information of genome is not available for plasmid profiling, the strains CB43 and ZQ1 used in this study were less than ten passages. Mice were sacrificed at 21 days post-infection, the inoculation site of the skin, the ankle joints, ears, and bladder were collected to quantitatively evaluate levels of colonization during infection as described in the section “Determination of Lyme borreliae burdens in infected ticks, tissues and blood samples.”.

### Generation of ticks carrying Lyme borreliae

The procedure of the tick infection has been described previously (21, 87). Basically, four-week-old male and female C3^−/−^ mice in BALB/c background were infected with 10^6^ of *B. afzelii* CB43, *B. garinii* ZQ1, or *B. burgdorferi* B31-5A4, B31-5A15, B31-5A4NP1Δ*cspA*-V or this *cspA* mutant strain producing CspA_B31_, CspA_PKo_, CspA_ZQ1_, or CspA_B31_L246D by intradermal injection as described above. The ear punches from those mice were collected at 13 days post infection, and DNA was extracted to perform qPCR using *Borrelia* 16S rRNA primers as previously described (26) (Table S2) to confirm the infection (See section “Determination of Lyme borreliae burdens in infected ticks, tissues and blood samples.”). At 14 days post infection, the uninfected larvae were allowed to feed to repletion on those spirochete-infected mice as described previously (21, 87). Approximately 100 to 200 larvae were allowed to feed on each mouse. The engorged larvae were collected and allowed to molt into nymphs in a desiccator at room temperature and 95% relative humidity in a room with light dark control (light to dark, 16:8 h). (21, 85).

### Serum resistance assays

*Coturnix* quail were subcutaneously injected with OmCI (1 mg/kg of quail) or PBS buffer, and the sera were collected at 6, 24, 48, 72, and 96 h post injection. A serum sensitive, high passaged *B. burgdorferi* strain B313 was cultivated to mid-log phase, followed by being diluted to a final concentration of 5×10^6^ bacteria/ml in BSKII medium without rabbit serum. These bacteria were then incubated with each of these quail serum samples (final concentration: 40% of serum). We also included the bacteria mixed with heat inactivated serum samples, which have been incubated at 56ºC for 2 h prior to being mixed with spirochetes. An aliquot was taken from each reaction at 0 and 4 h post injection to determine the number of motile bacteria under a Nikon Eclipse E600 darkfield microscope, as previously described (21). The survival percentage for those motile spirochetes was calculated using the number of mobile spirochetes at 4 h post incubation normalized to that at the very beginning of incubation with serum.

### Mouse, quail, and *P. leucopus* infection by ticks

The flat nymphs were placed in a chamber on four- to six-week old male and female BALB/c or C3^−/−^ mice in BALB/c background, and the engorged nymphs were collected from the chambers at seven days post nymph feeding as described (88). For ticks feeding on quail, the feathers located on the back of quail’s neck were plucked to expose approximately 2 to 3 cm^2^ of skin, close to the back of its head. 1.2 mL screw-top ‘cryo’ microcentrifuge vials (ThermoFisher Scientific) were cut to be used as mini chambers. The top of the caps from the chambers was pierced with a 25-gauge needle to create air holes, and sand papers were used to smooth any sharp edges along the cut surface edge of these chambers. Vetbond Tissue Adhesive (3M) was used to attach the chambers onto the exposed quail skin followed by manually restraining quail while the surgical glue dries (1-2 min). Ten nymphs were placed into the chambers on mice or quail, which those ticks to feed on. For OmCI-treated quail, the quail were subcutaneously injected with OmCI (1 mg/kg of quail) a day prior to the nymph feeding. The engorged nymphs were obtained from the chambers. The mice and quail were placed into a small cage, which then placed above the moat in a larger cage (for mice) or plastic bin (for quail). Ticks feeding on *Peromyscus leucopus* mice have been described previously (89, 90). Ten nymphs were placed in the ears of each mouse, five nymphs per ear and were allowed to feed until repletion. *P. leucopus* mice were separately placed in water cages which consisted of the cage being filled with approximately 2.5 cm of water along the bottom, a wire rack to keep the mouse out of the water, and then being placed in a larger hamster cage with water to prevent ticks from escaping. The engorged nymphs were recovered from the water cage beginning five days post nymph feeding. Blood and tick placement site from quail, mice, and *P. leucopus* mice were collected at seven days post nymph feeding.

### Feeding chamber assays by ticks

Artificial feeding chambers were prepared as described in our previous study (21). In short, the silicone rubber-saturated rayon membrane was generated as described (21), with the exception of adhering fiberglass mesh (3-mm pore; Lowe’s Inc.) to the membrane before attaching it to the rest of the chamber. Such membrane was attached to one side of a 2-cm length of polycarbonate tubing (hereafter called the chamber; inner diameter: 2.5 cm; outer diameter: 3.2 cm; (Amazon Inc.), as described, with the exception of using a rubber band to hold the chamber in place instead of a rubber O-ring (21). Feeding stimuli including hair and hair extract from white-tailed deer (*Odocoileus virginianus*) and a plastic tile spacer (Lowe’s Inc.) were added as described with the exception of using 3 stainless steel bearings (Amazon Inc.) instead of a nickel coin (21). *I. scapularis* nymphs carrying *B. burgdorferi* B31-5A4, *B. afzelii* CB43, *B. garinii* ZQ1; or *B. burgdorferi* B31-5A4, B31-5A15, B31-5A4NP1Δ*cspA*-V or this *cspA* mutant strain producing CspA_B31_, CspA_PKo_, CspA_ZQ1_, or CspA_B31_L246D were then added onto the chamber (5-8 ticks/chamber). Chamber feedings were carried out as previously described using human (BioIVT, Westbury, NY) or quail blood (Canola Poultry Market, Brooklyn, NY) (Fig. 2A). Chambers in blood were placed in a sealed Styrofoam cooler with wet paper towels to maintain humidity at approximately 87 to 95%. Depending on the experiments, blood was treated with CVF (ComTech) or OmCI to a final concentration of 17 μg/ml. Blood was changed daily, and was collected along with ticks after 5 days of feeding. SYBR-based qPCR was used to determine bacterial burdens in the ticks and blood using *Borrelia* 16S rRNA gene primers (Table S2).

### Determination of Lyme borreliae burdens in infected ticks, tissues and blood samples

The ticks fed on quail, mice, *P. leucopus* mice, or feeding chambers were homogenized by hand in a 1.5 ml Eppendorf tube (Eppendorf) with a plastic pestle (ThermoFisher Scientific). The DNA from tissues or blood or homogenized ticks was extracted using the EZ-10 Genomic DNA kit (Biobasic). The quantity and quality of DNA was assessed using a Nanodrop 1000 UV/Vis spectrophotometer (ThermoFisher Scientific). The 280:260 ratio was between 1.75 and 1.85, indicating the lack of contaminating RNA or proteins. qPCR was then performed to quantitate bacterial loads. Spirochete genomic equivalents were calculated using an ABI 7500 Real-Time PCR System (ThermoFisher Scientific) in conjunction with PowerUp SYBR Green Master Mix (ThermoFisher Scientific), based on amplification of the Lyme borreliae 16S rRNA gene using primers 16SrRNAfp and 16SrRNArp (Table S2), as described previously (23, 68). Cycling parameters for SYBR green-based reactions were 50°C for 2 min, 95°C for 10 min, and 45 cycles of 95°C for 15 s, 55°C for 30 s, and 60°C for 1 min. The number of 16S rRNA copies was calculated by establishing a threshold cycle (Cq) standard curve of a known number of 16S rRNA gene extracted from *B. burgdorferi* strain B31-5A4, then comparing the Cq values of the experimental samples.

### Sequence analysis of PFam54-IV

Nucleotide sequences of the PFam54-IV genes, including *bba68* (*cspA*_*B31*_), and *bba69* from *B. burgdorferi* B31 (AE000790), *pkoa0062*, *pkoa0063*, *pkoa0064*, *pkoa0065*, *pkoa0066,* and *pkoa0067* (*cspA*_*PKo*_), from *B. afzelii* PKo (CP002950), and *zqa67*, *zqa68* (*cspA_ZQ1_*), *zsa69*, *zsa70*, *zsa71*, and *zsa72* from *B. garinii* ZQ1 (AJ786369), were used as queries against the NCBI GenBank nr database using BLASTN(91). Sequences from organisms other than *B. burgdorferi*, *B. afzelii*, and *B. garinii* and duplicate sequences were discarded. The remaining sequences were then adjusted to include the full open reading frames with removal of any sequences containing premature stop codons. Sequences were grouped by species, and the sequences of the genes encoding the third and fifth clade of PFam54 (PFam54-III and Pfam54-V, respectively) from *B. burgdorferi* B31, *B. afzelii* PKo, and *B. garinii* ZQ1 were added to the sequence set (Genbank accession codes are found on on Fig. S5 to S7). Codon alignments were generated using T-Coffee on the TranslatorX server (92, 93), followed by calculation of pairwise identity in Clustal Omega (94). Only Pfam54-IV best hits were kept. The remaining sequences were then realigned by codon and pairwise identity was calculated as above. The percent identity between each of these genes and each of the identified PFam54-IV alleles is shown in Fig. S5 to S7.

### Generation of recombinant PFam54 proteins

The pQE30Xa vectors encoding the open reading frames lacking the putative signal sequences of *bba68* (*cspA*_*B31*_), *bba69* from *B. burgdorferi* strain B31-5A4, *mmsa67*, *mmsa68*, *mmsa69*, or *mmsa70*, and *mmsa71* (*cspA*_*MMS*_) from *B. afzelii* strain MMS, or *zqa67*, *zqa68* (*cspA_ZQ1_*), *zsa69*, *zsa70*, *zsa71*, or *zsa72* from *B. garinii* strain ZQ1 were obtained previously to generate recombinant histidine-tagged proteins (24). The plasmids were transformed into *E. coli* strain M15, and the plasmid inserts were sequenced using Sanger sequencing on an ABI 3730xl DNA Analyzer (ThermoFisher Scientific) at the NYSDOH Wadsworth Center ATGC Core Facility. The resulting M15 derived strains were used to produce respective recombinant PFam54-IV (Table S1). The histidine-tagged PFam54-IV were produced and purified by nickel affinity chromatography with Ni-NTA agarose according to the manufacturer’s instructions (Qiagen, Valencia, CA).

### FH binding assay by qualitative ELISA

Qualitative ELISA for FH binding by PFam54-IV was performed as described (21, 95). One microgram of BSA (negative control; Sigma-Aldrich, St. Louis, MO) or FH from mouse or quail was coated onto microtiter plate wells by incubating the plate for overnight at 4ºC. Then, 100 μl 2 μM of histidine-tagged DbpA from *B. burgdorferi* strain B31 (negative control) (96) or each of the PFam54-IV was added to the wells. Mouse anti-histidine tag 1:200× (Sigma-Aldrich, St. Louis, MO) and HRP-conjugated goat anti-mouse IgG 1:1,000× (Seracare Life Sci., Inc, Milford, MA) were used as primary and secondary antibodies, respectively, to detect the binding of histidine-tagged proteins. The plates were washed three times with PBST (0.05% Tween 20 in PBS), and 100 μl of tetramethyl benzidine (TMB) solution (ThermoFisher Scientific) was added to each well and incubated for 5 min. The reaction was stopped by adding 100 μl of 0.5% hydrosulfuric acid to each well. Plates were read at 405 nm using a Tecan Sunrise Microplate reader at five minutes after the incubation (Tecan Life science, Männedorf, Switzerland).

### Phylogenetic reconstruction

The PFam54-IV codon alignment from *B. burgdorferi* B31, *B. afzelii* MMS, and *B. garinii* ZQ1 was used to generate a Bayesian phylogenetic reconstruction was carried out in BEAST v1.8.4 with a relaxed lognormal clock, an estimated mutation rate of 4.75 x 10^−6^ substitutions/site/year and a coalescent Bayesian skyline model (33, 97). A Markov chain Monte Carlo chain length of 10,000,000 steps was used with a 100,000-step thinning, resulting in effective sample sizes greater than 200, an indication of an adequate chain mixing. The resulting maximum clade credibility tree was visualized in FigTree v1.4.4 (98). We evaluated alternative, competing evolutionary scenarios for PFam54-IV based on species-specific divergence, or clustering by FH-binding activity shown in Fig. S10A (99). A battery of statistical phylogenetic tests was deployed in IQ-TREE (Kishino-Hasegawa, Shimodaira-Hasegawa, Expected Likelihood Weight, and Approximately Unbiased tests)(101–104).

### Statistical analysis

Significant differences between samples were assessed using the Mann-Whitney *U* test or the Kruskal-Wallis test with the two-stage step-up method of Benjamini, Krieger, and Yekutieli. A P-value < 0.05 (*) or (^#^) was considered to be significant (105).

## ACKNOWLEDGEMENTS

We thank Frank Blaisdell, and Dierdre Torrisi from the Wadsworth vet sciences facility and Ashley Marcinkiewicz and Patricia Lederman for animal husbandry, Levi Poirier and Ing-Nang Wang for assistance with gene sequencing and phylogenetic reconstructions, Ashley Marcinkiewicz for critical reading of the manuscript, Patricia Rosa for sharing the unpublished observation of *B. afzelii* strain PKo, and Roxie Giradin for assistance with SDS-PAGE. We thank Wadsworth ATGC core for plasmid sequencing, Leslie Eisele of Wadsworth Biochemistry and Immunology Core for HPLC performance, and Karen Chave of the Wadsworth Protein Expression Core for purifying factor H. This work was supported by NIH-U01CK000509 (DMT and MDW), NSF-IOS1755286 (DMT, MDW, SOK, ADII, LDK, TMH, and YL), DoD-TB170111, NIH-R21AI144891, NIH-R21AI146381, New York State Department of Health Wadsworth Center Start-Up Grant (TMH and YL), the Czech Science Foundation grant No. 17-21244S (ROMR), NIH R01AI121401 (PK), and the LOEWE Center DRUID Novel Drug Targets against Poverty-Related and Neglected Tropical Infectious Diseases, project C3 (PK). The funders had no role in study design, data collection and analysis, decision to publish, or preparation of the manuscript. The authors declare that the research was conducted in the absence of any commercial or financial relationships that could be construed as a potential conflict of interest.

