## Supplemental Text for "Host tropism determination by convergent evolution of immunological evasion in the Lyme disease system"

**SUPPLEMENTARY INFORMATION**

**SUPPLEMTARY FIGURE LEGENDS**

**Figure S1. Lyme borreliae vary in their ability to infect mice via intradermal injection of BALB/c but not C3^-/-^ BALB/c mice.** **(A to D)** BALB/c or **(E to H)** C3^-/-^ BALB/c mice were injected with 10^6^ *B. burgdorferi* B31-5A4 (“*Bb* B31-5A4”), *B. garinii* ZQ1 (“*Bg* ZQ1”), or *B. afzelii* CB43 (“*Ba* CB43”). At 21 days post injection (“dpi”), spirochete burdens were determined in the **(A and E)** inoculation site (“Inoc. site”), **(B and F)** ears, **(C and G)** bladder, **(D and H)** ankles using qPCR and normalized to 100 ng total DNA. For C3^-/-^ BALB/c mice, uninfected *I. scapularis* larvae were placed on those mice at 14 days post injection (“dpi”) and allowed to feed to repletion, and some of the replete larvae were permitted to molt to flat nymphs. Spirochete burdens in **(I)** replete larvae, and **(J)** post molting flat nymphs (“flat nymphs”) were determined using qPCR. The above-mentioned tissues from uninfected mice and uninfected nymphs were included as control (“Uninfect.”). For the burdens in tissue samples, the resulting values were normalized to 100ng total DNA in those tissues. Shown are the geometric means of bacterial loads ± 95% confidence interval of bacterial burdens from 6 replete larvae, flat nymphs or tissues from 5 BALB/c mice or indicated numbers of C3^-/-^ BALB/c mice (9 *Bb* B31-5A4-infected inoculation sites and ears, 6 *Bg* ZQ1-infected inoculation sites and ears, or 5 all other tissues). Significant differences (p < 0.05, Kruskal-Wallis test with Two-stage step-up method of Benjamini, Krieger, and Yekutieli) in the spirochete burdens relative to uninfected ticks or tissues (‘‘*’’) are indicated.

**Figure S2. The complement of quail inoculated with OmCI is inactivated.** *Coturnix* quail were subcutaneously injected with OmCI (1 mg/kg of quail) or PBS buffer. Untreated (black bars) or heat-treated (white bars) of sera collected from these quail at indicated time points after inoculation were incubated with a serum sensitive, high passaged *B. burgdorferi* strain B313 for 0- and 4-h with a final concentration of 40%. The number of motile spirochetes was assessed microscopically. The survival percentage of the spirochetes was calculated using the number of mobile spirochetes at 4-h post incubation normalized to that at 0-h of incubation with serum. Each bar represents the mean of three independent determinations ± SEM from sera from five quail per group. Significant differences (P < 0.05 by Kruskal-Wallis test with the two-stage step-up method of Benjamini, Krieger, and Yekutieli) in the percentage survival of spirochetes incubated with untreated sera from OmCI-inoculated quail, compared to that in heat inactivated sera from those quail (‘‘*’’).

**Figure S3. Protein sequence alignment of CspA variants from *B. afzelii* CB43, PKo, and MMS and the synteny from PFam54-IV proteins of these strains.** **(A)** Amino acid alignments of CspA variants from *B. afzelii* CB43, PKo, and MMS aligned in M-Coffee. Blue shading indicates amino acids conserved among the three variants. **(B)** Synteny of Pfam54-IV proteins from *B. afzelii* CB43, PKo, and MMS. One-to-one orthologs are colored accordingly. Note that *pkoa0066* is only present in the strain PKo but not others. Figure is drawn to scale; scale bar denotes 500 bp.

**Figure S4. CspA variants confer differential transmissibility from nymphs to *P. leucopus* mice.** *I. scapularis* nymphs infected by WT *B. burgdorferi* B31-5A15 (“B31-5A15”), *B. burgdorferi* B31-5A4NP1*ΔcspA* transformed with an empty shuttle vector (“Vector”), or this deletion strain producing a mutated variant of CspA from *B. burgdorferi* B31 selectively devoid of FH binding activity (“L246D”), WT *B. burgdorferi* B31 (“B31”), *B. garinii* ZQ1 (“ZQ1”), or *B. afzelii* PKo (“PKo”) were allowed to feed to repletion on *P. leucopus* mice. Uninfected nymphs and *P. leucopus* mouse tissues and blood were included as control (“Uninfect.”). Fed nymphs were collected upon repletion, and blood and tissues were collected at 7 days post nymph feeding (“dpf”). Spirochete burdens in **(A)** replete nymphs, **(B)** tick bite sites of skin (“Inoc. site”), and **(C)** blood were determined by qPCR. For the burdens in tissue samples, the resulting values were normalized to 100ng of total DNA. Shown are the geometric means of bacterial loads ± 95% confidence interval of bacterial burdens in tissues from five *P. leucopus* mice per group or replete nymphs (8 nymphs carrying the strain B31-5A15, 12 nymphs carrying the strain “Vector”, 13 nymphs carrying the strain pCspA-B31, 15 nymphs carrying the strain pCspA-PKo, 13 nymphs carrying the strain pCspA-ZQ1, or 10 nymphs carrying the strain pCspA-L246D). Significant differences (P < 0.05, Kruskal-Wallis test with the two-stage step-up method of Benjamini, Krieger, and Yekutieli) in the spirochete burdens relative to uninfected ticks or *P. leucopus* mouse tissues or blood (‘‘*’’) are indicated.

**Figure S5. Pairwise comparisons of codon alignments define CspA variants in *B. burgdorferi.*** **(A)** Nucleotide sequences encoding PFam54-IV proteins from *B. burgdorferi* B31 were used as queries to mine NCBI GenBank for orthologs. **(B)** Sequence identity values of pairwise comparisons were plotted against the number of comparisons with those respective values (bin width: 0.25). Clear breaks in the pairwise sequence identity distribution were inspected to differentiate highly identical (> 95%) from moderately divergent comparisons (< 80%). The Pfam54-IV ortholog from a particular *B. burgdorferi* strain with more than 95% identity to CspA_B31_ was defined as the CspA orthologs in that strain.

**Figure S6. Pairwise comparisons of codon alignments define CspA variants in *B. afzelii.* (A)** Nucleotide sequences encoding PFam54-IV proteins from *B. afzelii* PKo were used as queries to mine NCBI GenBank for orthologs. **(B)** The identity values of comparisons were plotted versus the number of comparisons with those respective values, with bin widths of 0.25. Clear breaks in the pairwise sequence identity distribution were inspected to differentiate the highly identical (> 98%) from moderately divergent comparisons (< 80%). The Pfam54-IV ortholog from a particular *B. afzelii* strain with more than 98% identity to CspA_B31_ was defined as the CspA orthologs in that strain.

**Figure S7. Pairwise comparisons of codon alignments define CspA variants in *B. garinii.*** **(A)** Nucleotide sequences encoding PFam54-IV proteins from *B. garinii* ZQ1 were used as queries to mine NCBI GenBank for orthologs. **(B)** The identity values of comparisons were plotted versus the number of comparisons with those respective values, with bin widths of 0.25. Clear breaks in the pairwise sequence identity distribution were inspected to differentiate the highly identical (> 93%) from moderately divergent comparisons (< 80%). The Pfam54-IV ortholog from a particular *B. garinii* strain with more than 93% identity to CspA_B31_ was defined as the CspA orthologs in that strain.

**Figure S8. Phylogenetic reconstruction of PFam54-IV genes from the strains of *B. burgdorferi*, *B. afzelii*, and *B. garinii* available from GenBank.** Maximum likelihood phylogenetic reconstruction was generated based on codon alignments of the genes encoding Pfam54-IV from the strains of *B. burgdorferi*, *B. afzelii*, and *B. garinii* available on GenBank. Sequences were aligned by codon using T-Coffee on the TranslatorX sequence alignment server. Phylogenetic reconstructions were generated based on the resulting codon alignment using IQ-TREE with model finder, 1,000 ultrafast bootstrap replicates, and 1,000 adjusted likelihood ratio test replicates. CspA variants of *B. burgdorferi*, *B. afzelii*, and *B. garinii* are colored in blue, green, and orange, respectively.

**Figure S9. Competing evolutionary scenario evaluation of PFam54-IV gene divergence rejects alternative tree topologies of PFam54-IV proteins forming clades by FH-binding activities or spirochete genospecies. (A)** Phylogenetic scenarios showing a single emergence of FH-binding or PFam54-IV genes forming clades by species. **(B)** The log-likelihood for each tree (logL), the RELL bootstrap proportion (bp-RELL; 1 is total support for that tree), Kishino-Hasegawa test (p-KH; 1 supports the tree, <0.05 rejects), Shimodaira-Hasegawa test (p-SH; 1 supports the tree, <0.05 rejects), Expected Likelihood Weight test (c-ELW; 1 is total support for that tree), and the Approximately Unbiased test (p-AU; 1 supports the tree, <0.05 rejects).

**SUPPLEMENTAL TABLES**

**Table S1. Bacteria strains, and plasmids used in this study**

| Strain or plasmid | Genotype or characteristic | Sources |
| --- | --- | --- |
| *Borrelia* strains |  |  |
| ZQ1 | Clonal isolate of *B. garinii* strain ZQ1 | (106) |
| PKo | Clonal isolate of *B. afzelii* strain PKo | (25) |
| CB43 | Clonal isolate of *B. afzelii* strain CB43 | (107) |
| B31-5A15 | Clonal isolate of *B. burgdorferi* strain B31 lacking lp21 | (85) |
| B31-5A4NP1Δ*cspA* | B31-5A4NP1, the clonal isolate of strain B31 with *bbe02*:: KanR^a^, *cspA*::StrR^b^ , and lacking lp21 | (108) |
| B31-5A4NP1Δ*cspA* /pBSV2G | B31-5A4NP1Δ*cspA* carrying plasmid pBSV2G | (21) |
| B31-5A4NP1Δ*cspA* /pBSV2G-CspA_B31_ | B31-5A4NP1Δ*cspA* complemented with intact *cspA* (*bba68*) from *B. burgdorferi* strain B31 under the control of *cspA* promoter from this strain (PcspA). | (21) |
| B31-5A4NP1Δ*cspA* /pBSV2G-CspA_PKo_ | B31-5A4NP1Δ*cspA* complemented with intact *cspA* (*bafPKo_A0067*) from *B. afzelii* strain PKo under the control of PcspA. | (21) |
| B31-5A4NP1Δ*cspA* /pBSV2G-CspA_ZQ1_ | B31-5A4NP1Δ*cspA* complemented with intact *cspA* (*zqa68*) from *B. garinii* strain ZQ1 under the control of PcspA. | (21) |
| B31-5A4NP1Δ*cspA* /pBSV2G-CspA_B31_L246D | B31-5A4NP1Δ*cspA* complemented with intact *cspA* from *B. burgdorferi* strain B31 with leucine-246 replaced by aspartate under the control of PcspA. | (21) |
| *E. coli* strains |  |  |
| DH5α | F- Φ80lacZΔM15 Δ(lacZYA-argF) U169 recA1 endA1 hsdR17(rk-, mk+) phoA supE44 thi-1 gyrA96 relA1 λ- | ThermoFisher |
| BL21 | F–, ompT, hsdSB (rB–, mB–), dcm, gal, λ(DE3) | Promega |
| M15 [Prep4] | F-, Φ80ΔlacM15, thi, lac-, mtl-, recA+ , KmR | Qiagen |
| BL21/pET30a-DbpA_B31_ | BL21 producing histidine tagged residue 26 to 192 of DbpA from *B. burgdorferi* strain B31 | (96) |
| M15 [Prep4]/pQE-CspA_B31_ | M15 producing histidine tagged residue 26 to 252 of CspA (BBA68) from *B. burgdorferi* strain B31 | (21) |
| M15 [Prep4]/pQE-BBA69 | M15 producing histidine tagged residue 26 to 264 of BBA69 from *B. burgdorferi* strain B31 | This study |
| M15 [Prep4]/pQE- MMSA67 | M15 producing histidine tagged residue 20 to 232 of MMA67 from *B. afzelii* strain MMS | This study |
| M15 [Prep4]/pQE- MMSA68 | M15 producing histidine tagged residue 20 to 240 of MMSA68 from *B. afzelii* strain MMS | This study |
| M15 [Prep4]/pQE- MMSA69 | M15 producing histidine tagged residue 23 to 269 of MMSA69 from *B. afzelii* strain MMS | This study |
| M15 [Prep4]/pQE- MMSA70 | M15 producing histidine tagged residue 26 to 239 of MMSA70 from *B. afzelii* strain MMS | This study |
| M15 [Prep4]/pQE-CspA_MMS_ | M15 producing histidine tagged residue 28 to 242 of CspA (MMSA71) from *B. afzelii* strain MMS | (24) |
| M15 [Prep4]/pQE-ZQA67 | M15 producing histidine tagged residue 26 to 251 of ZQA67 from *B. garinii* strain ZQ1 | This study |
| M15 [Prep4]/pQE-CspA_ZQ1_ | M15 producing histidine tagged residue 27 to 256 of CspA (ZQA68) from *B. garinii* strain ZQ1 | (24) |
| M15 [Prep4]/pQE-ZSA69 | M15 producing histidine tagged residue 26 to 236 of ZSA69 from *B. garinii* strain ZQ1 | This study |
| M15 [Prep4]/pQE-ZSA70 | M15 producing histidine tagged residue 26 to 249 of ZSA70 from *B. garinii* strain ZQ1 | This study |
| M15 [Prep4]/pQE-ZSA71 | M15 producing histidine tagged residue 26 to 239 of ZSA71 from *B. garinii* strain ZQ1 | This study |
| M15 [Prep4]/pQE-ZSA72 | M15 producing histidine tagged residue 25 to 257 of ZSA72 from *B. garinii* strain ZQ1 | This study |
| Plasmid |  |  |
| pJET1.2/Blunt | AmpR^c^; PCR cloning vector | ThermoFisher |
| pQE30Xa | AmpR^c^; histidine-tag protein expression vector | Qiagen |
| pQE-CspA_B31_ | pQE30Xa encoding histidine fusion protein residue 26 to 252 of CspA (BBA68) from *B. burgdorferi* strain B31 | (21) |
| pQE-BBA69 | pQE30Xa encoding histidine fusion protein residue 26 to 264 of BBA69 from *B. burgdorferi* strain B31 | This study |
| pQE-MMSA67 | pQE30Xa encoding histidine fusion protein residue 20 to 232 of MMA67 from *B. afzelii* strain MMS | This study |
| pQE-MMSA68 | pQE30Xa encoding histidine fusion protein residue 20 to 240 of MMSA68 from *B. afzelii* strain MMS | This study |
| pQE-MMSA69 | pQE30Xa encoding histidine fusion protein residue 23 to 269 of MMSA69 from *B. afzelii* strain MMS | This study |
| pQE-MMSA70 | pQE30Xa encoding histidine fusion protein residue 26 to 239 of MMSA70 from *B. afzelii* strain MMS | This study |
| pQE-CspA_MMS_ | pQE30Xa encoding histidine fusion protein residue 28 to 242 of CspA (MMSA71) from *B. afzelii* strain MMS | This study |
| pQE-ZQA67 | pQE30Xa encoding histidine tagged protein residue 26 to 251 of ZQA67 from *B. garinii* strain ZQ1 | This study |
| pQE-CspA_ZQ1_ | pQE30Xa encoding histidine tagged protein residue 27 to 256 of CspA (ZQA68) from *B. garinii* strain ZQ1 | (24) |
| pQE-ZSA69 | pQE30Xa encoding histidine tagged protein residue 26 to 236 of ZSA69 from *B. garinii* strain ZQ1 | This study |
| pQE-ZSA70 | pQE30Xa encoding histidine tagged protein residue 26 to 249 of ZSA70 of Zsa70 from *B. garinii* strain ZQ1 | This study |
| pQE-ZSA71 | pQE30Xa encoding histidine tagged protein residue 26 to 239 of ZSA71 from *B. garinii* strain ZQ1 | This study |
| pQE-ZSA72 | pQE30Xa encoding histidine tagged protein residue 25 to 257 of ZSA72 of Zsa72 from *B. garinii* strain ZQ1 | This study |
| pQE-CspA_B31_L246D | pQE30Xa encoding histidine fusion protein residue 26 to 251 of CspA (BBA68) from *B. burgdorferi* strain B31 with leucine-246 replaced by aspartate | (21) |
| pBSV2G | GenR^d^; pBSV2-derived shuttle vector. | (109) |
| pBSV2G-CspA_B31_ | pBSV2G encoding intact *cspA* (*bba68*) from *B. burgdorferi* strain B31 under the control of *cspA* promoter from this strain (PcspA). | (21) |
| pBSV2G-CspA_PKo_ | pBSV2G encoding intact *cspA* (*bafPKo_A0067*) from *B. afzelii* strain PKo under the control of pCspA | (21) |
| pBSV2G-CspA_ZQ1_ | pBSV2G encoding intact *cspA* (*zqa68*) from *B. garinii* strain ZQ1 under the control of pCspA | (21) |
| pBSV2G-CspA_B31_L246D | pBSV2G encoding intact *cspA* (*bba68*) from *B. burgdorferi* strain B31 under the control of pCspA with leucine-246 replaced by aspartate | (21) |

KanR^a^, Kanamycin resistant

StrR^b^, Streptomycin resistant

AmpR^c^, Ampicillin resistant

GenR^d^, Gentamicin resistant

**Table S2. Primers used in this study.**

| Primer/Vector | Sequence | Amplified DNA fragment |
| --- | --- | --- |
| 16srRNAfp | GCTTCGCTTGTAGATGAGTCTGC | *BB16srRNA* |
| 16srRNArp | TTCCAGTGTGACCGTTCACC |  |
| ColE1fp | CTACATACCTCGCTCTGCTAATC | *BBcolE1* |
| ColE1rp | CGAAACCCGACAGGACTATAAA |  |
| mNidfp | CCAGCCACAGAATCCCATCC | *mNidogen* |
| mNidrp | GGACATACTCTGCTGCCATC |  |
| Qβ-actinfp | CTGGCACCTAGCACAATGAA | *qβ-actin* |
| Qβ-actinrp | CTGCTTGCTGATCCACATCT |  |
| Kanfp | ATGAGCCATATTCAACGGGAA | Kanamycin |
| Kanrp | TTAGAAAAACTCATCGAGCAT |  |
| Genfp | ATGTTACGCAGCAGCAAC | Gentamycin |
| Genrp | TTAGGTGGCGGTACTTGG |  |
| Strfp | CAGGATGACGCCTAACAA | Streptomycin |
| Strrp | CCACCTTCAACAGATCGC |  |
